# Geographic and Climatic Origins Shape the Leaf Metabolome of *Populus trichocarpa*

**DOI:** 10.64898/2026.05.20.726516

**Authors:** Moritz Popp, Sol Yepes-Vivas, Ina Zimmer, Athena McKown, Charles Hefer, Basem Kanawati, Philippe Schmitt-Kopplin, Shawn D. Mansfield, Sybille B. Unsicker, Jürgen Elthing, Jörg-Peter Schnitzler

**Affiliations:** Research Unit Environmental Simulation (EUS), Helmholtz Zentrum München, 85764 Neuherberg, Germany; Plant-Environment-Interactions Group, Botanical Institute, Kiel University, 24118 Kiel. Germany; Department of Forest and Conservation Sciences, Faculty of Forestry, University of British Columbia, Vancouver, British Columbia, V6T 1Z4, Canada; Department of Botany, Faculty of Science, University of British Columbia, Vancouver, British Columbia, V6T 1Z4, Canada; Research Unit Analytical BioGeoChemistry, Helmholtz Zentrum München, 85764, Neuherberg, Germany; Department of Wood Science, Faculty of Forestry, University of British Columbia, Vancouver, British Columbia, V6T 1Z4, Canada; University of Victoria, Centre for Forest Biology & Department of Biology, Victoria BC, Canada; Ecosystem Physiology, Faculty of Environment and Natural Resources, University Freiburg, D-79110 Freiburg, Germany

**Keywords:** chemodiversity, FT-ICR-MS, LC-MS/MS, Functional Hill Diversity, isoprene, metabolomics, *Populus trichocarpa*, provenance, specialised metabolism

## Abstract

- Background and Aims: Chemodiversity is a fitness-relevant trait shaped by genetics, environment, and their interaction. *Populus trichocarpa* naturally inhabits broad climatic gradients and shows extensive variation in specialised metabolism. We investigated whether provenance and climate of origin imprint leaf chemodiversity and class-level relationships under common-garden conditions, and how these patterns relate to gene expression.
- Methods: Leaves from 87 *P. trichocarpa* genotypes representing 22 provenances from the west coast of North America growing in a common garden were profiled by untargeted FT-ICR-MS (1030 features) and targeted LC-MS/MS. A subset of 41 genotypes was subject to RNA-seq analyses. We tested whether provenance influenced multivariate patterns and whether metabolomic differences were related to geographic and climatic distance, where chemodiversity was quantified as Functional Hill Diversity.
- Key Results: *P. trichocarpa* metabolomes differed among origins despite shared growth conditions and showed distance-decay with both geography and climate. North-south extremes were well separated, and within-drainage samples shared high similarity. Flavonoid and isoprenoid pools strongly co-varied across individuals, whereas isoprene synthase activity did not predict total isoprenoids. Transcriptomes showed within-pathway coherence but limited overall provenance separation.
- Conclusions: Leaf chemistry in *P. trichocarpa* retains signatures of geographic origin even under common-garden conditions. Coordinated investment in flavonoids and isoprenoids, together with among-origin differences in functional chemodiversity, reveals provenance-linked chemical fingerprints that complement genomic and metabolic trait data for climate-informed deployment.

## Introduction

Plant secondary metabolites, also known as specialised metabolites, are extraordinarily diverse. To date, more than 130,000 natural compounds have been catalogued in plants alone, representing around 70% of all known natural products (Ntie-Kang & Svozil, 2020). Nearly three-quarters of known natural plant products are non-nitrogen-containing compounds, including terpenoids (products containing C5H8 isoprene units) and phenolics (spanning simple aromatic compounds, phenylpropanoid esters and amides, flavonoids, and lignans). Nitrogen-containing alkaloids constitute another major group. Chemical diversity varies not only among lineages and species, but also within species, consistent with local adaptation across populations.

Plant phenotypic traits are shaped by genetic and environmental factors, as well as by their interaction (Nicotra et al., 2010). Understanding the drivers of phenotypic variation, including phenotypic plasticity, may help predict the capacity of plant species to persist under unfamiliar conditions, including those imposed by climate change. Gene-by-environment interactions and their effects on phenotypic variation are often examined in common gardens or controlled environments, typically focusing on individual stressors such as herbivory (Bertic et al., 2021) or drought (Kerr et al., 2023). In addition to physiological and morphological traits, many studies reveal extensive chemical diversity, with individual metabolites or groups of compounds associated with functional traits.

Secondary metabolites are derived from primary metabolites through diverse biosynthetic pathways and can mediate an array of biological functions, including defence against herbivores and pathogens (Bertic et al., 2021), attraction of pollinators (Ayelo et al., 2021; Derstine et al., 2020), protection against UV radiation (Brown et al., 2005), and plant-to-plant and plant-insect communication (Hagiwara et al., 2021; Meents & Mithöfer, 2020). These diverse bioactive compounds have also been explored extensively for pharmaceutical applications (Pal et al., 2021). The diversity of metabolites within a plant, tissue, or population is often termed ’chemodiversity’ and is increasingly recognised as a plant functional trait (Wetzel & Whitehead, 2020; Thon et al., 2024; Hanusch et al., 2026). Most such compounds arise from specialised metabolism, which is more flexible than primary metabolism and therefore offers greater variability in both structure and abundance. As a functional trait, chemodiversity can influence plant fitness (Müller & Junker, 2022).

In line with ecological methodology, chemodiversity can be quantified using diversity measures based on Hill numbers, which combine richness and evenness into a single metric (Chao et al., 2014). Similarity-sensitive extensions incorporate distances between entities, such that chemically similar metabolites contribute less than chemically distinct metabolites (Leinster & Cobbold, 2012). Distance-based functional diversity measures, grounded in Hill numbers, provide an explicit framework for this extension (Chiu & Chao, 2014). In metabolomics, these approaches have been used to quantify Functional Hill Diversity (Petrén et al., 2023), combining richness, evenness, and chemical similarity into a single index that captures abundance structure and chemical-space coverage.

Chemodiversity and its functional relevance are frequently investigated in chemically rich systems, including herbaceous plants and trees. Examples include *Tanacetum vulgare* (Clancy et al., 2018), *Solanum dulcamara* (Calf et al., 2018), and tree species such as oak (*Quercus robur*) and poplar (*Populus* x *canescens*) (Bertic et al., 2021; Yepes-Vivas et al., 2025). Poplar species are characterized by high diversity and pronounced variation in the abundance of secondary metabolites, particularly terpenoids and phenolics, including simple phenylpropanoids, flavonoids, and condensed tannins (Madritch & Lindroth, 2015). In poplar, metabolites produced for defence or in response to water availability are known to be influenced by genotype and environment (Barchet et al., 2014; Hamanishi et al., 2015; Rubert-Nason et al., 2015). Moreover, despite their distinct biochemical pathways, the production of entire classes of secondary metabolites can be linked. For example, genetically eliminating isoprene production in poplar reduces flavonoid levels (Behnke et al., 2010).

Here, we investigate chemodiversity and its ecological relevance in a chemically rich tree system. *Populus trichocarpa* Hook. (black cottonwood) spans extensive latitudinal and climatic gradients along the Pacific coast of North America, and common-garden studies show that multiple traits are associated with geography and climate of origin (McKown et al., 2014a; Porth et al., 2013), and genome-wide association approaches implicate many loci underlying ecological trait variations (McKown et al., 2014b). Despite these advances, the influence of range-wide provenance structure and climate at place of origin on leaf chemodiversity under shared growing conditions, and the relationship between provenance-linked chemical patterns and biosynthetic gene expression, remain poorly resolved.

In this study, we sampled developmentally similar leaves from 87 genotypes representing 22 *P. trichocarpa* provenances grown in a common garden and combined untargeted ultrahigh-resolution FT-ICR-MS with targeted LC-MS and RNA-seq (using a 41-tree subset). We focused on three aims: (i) analysing provenance and climate-of-origin signals in metabolomic composition and chemodiversity; (ii) testing for covariation between major metabolite classes (flavonoids and isoprenoids) and its relationship to isoprene synthase activity; and (iii) assessing whether transcript patterns in key biosynthetic pathways exhibit coherence consistent with metabolite-class relationships.

## Material and Methods

### Common garden design at the University of British Columbia, Vancouver, Canada

The common garden is located at Totem Field on the University of British Columbia, Canada (49°15’N, 123°14’W). Trees of *Populus trichocarpa* Hook. from the west coast of North America were transferred from common gardens in Surrey and Terrace, British Columbia, in 2008. The source common garden was established in 2000 using cuttings from wild trees collected by the British Columbia Ministry of Forests. The common garden at Totem Field was planted in a random block design with an initial spacing of 1.5 x 1.5 m. The field was watered regularly and mowed to suppress weed growth (McKown et al., 2014c). Tree samples were identified by a four-letter code followed by a number. The first three letters represent the river drainage from which the samples were originally collected (e.g. DEN for Dean River). The fourth letter identifies the specific location along the river, and the number is a unique identifier for each tree from that location (e.g. DENA 17.1). In total, 87 genotypes from 22 unique locations spanning 44° N to 55° N were sampled. The origins of the samples, together with the location of the common garden, are shown in Figure 1 (Fig. 1D). For each tree, from a south-facing, upper canopy branch, the first unfurled / fully flattened leaf (excluding the petiole) from the leading branch tip was sampled. Leaves from two distinct branches were collected as biological replicates, and each leaf was divided into two subsamples and frozen immediately at -80 °C. Sampling was carried out on a sunny day in August. One subsample was used for RNA-seq analysis, and the corresponding subsample was shipped to the Helmholtz Zentrum for isoprene synthase activity measurements, FT-ICR-MS, and LC-MS analyses.

**Fig 1.**
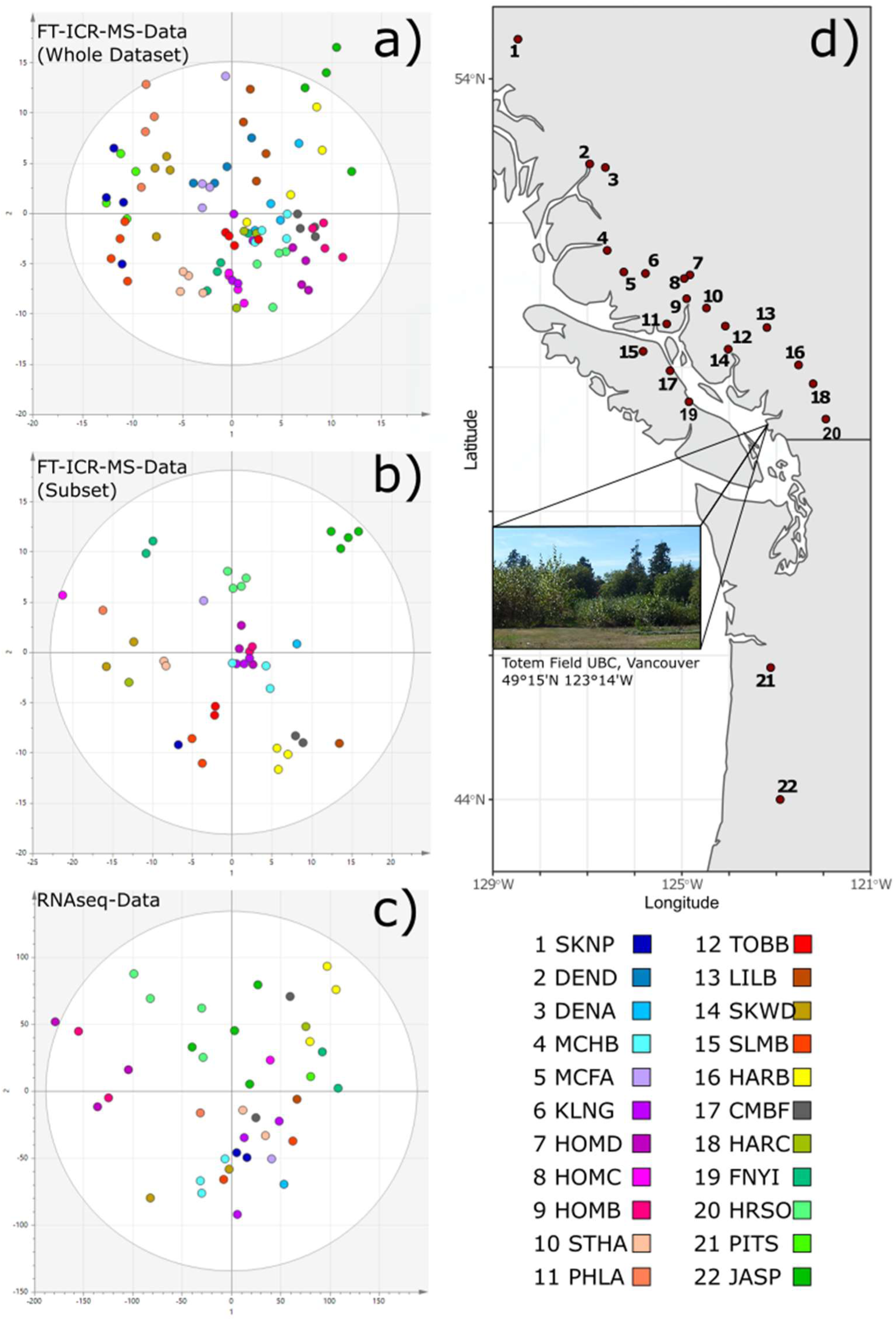
Multivariate separation of genotype provenances revealed by metabolomic and transcriptomic O2PLS-DA analyses. a: Orthogonal Two-Partial Least Squares Discriminant Analysis (O2PLS-DA) of a Fourier-transform ion cyclotron resonance mass spectrometry (FTICR-MS) metabolomic dataset from 87 individual genotypes (from 22 provenances / places of origin named and colour-coded in the bottom right), showing the separation across the first and second axis (R2X(cum) = 0.346, Q(cum) = -0.002). b) O2PLS-DA of the metabolomic dataset from those genotypes, showing the separation across the first and second axis (R2X(cum) = 0.241, Q(cum) = -0.020). c) O2PLS-DA of the RNAseq-data set for 41 trees showing the separation on the first and second axis (R2X(cum) = 0.413, Q(cum) = -0.037). Data points are colour-coded according to provenance in (a), (b), (c). (d) Geographic map of the west coast of North America showing the places of origins of the 22 different provenances, which were numbered by increasing latitude. The location of the common garden near Vancouver, BC is also indicated, and a photograph of the site is shown in the inset.

### Metabolomic analysis by Fourier transform ion cyclotron resonance mass spectrometry

For FT-ICR-MS analysis, frozen leaf material was ground to a fine powder in liquid nitrogen in a mortar and pestle. Twenty mg powder was then mixed with pre-cooled extraction solvent (methanol:isopropanol:water, 1:1:1), vortexed, and sonicated for 15 min in a cooled ultrasonic bath. Samples were then shaken for 10 min at 4 °C and centrifuged (4 °C, 14,000 rpm, 10 min). Supernatants were diluted 1:25 in brown glass vials. Each sample was analysed in duplicate.

Measurements were performed on a Fourier transform ion cyclotron resonance mass spectrometer (FT-ICR-MS; APEX Qe, Bruker, Bremen, Germany) equipped with a 12 T superconducting magnet and an APOLLO II ESI source, at a flow rate of 120 μL h^−1^ using a Gilson autosampler and a micro electrospray source (Kaling et al., 2015, 2018). Spectra were acquired in negative ion mode over 122–1000 m/z with 300 scans per acquisition. Ions were accumulated for 300 ms in the collision cell. The average resolving power was ∼400,000 at 400 m/z with a 4 MW transient size and transient length >1.6 s. Samples were measured in random order to minimise batch and memory effects.

After internal calibration, peak lists were exported with Bruker Data Analysis and processed in R. We filtered FT artefacts (side lobes/’wiggles’) using a resolution-based approach (Kanawati et al., 2017), removed isotopes and mass-defect artefacts, and aligned peaks across samples (1 ppm tolerance). Molecular sum formulas were assigned from exact mass, and candidate formulas were filtered using the ’Seven Golden Rules’ (Kind & Fiehn, 2007). Peaks not detected in at least 50% of samples were removed. Duplicate measurements were averaged and intensities were normalised to total ion current. The resulting m/z peak list was annotated with the MassBank of North America (MoNA) database using an error tolerance of 1 ppm and, in a second pass, with an in house-generated database containing m/z peaks annotated in earlier poplar FT-ICR-MS experiments (Kaling et al., 2015, 2018). In both cases, the best match (smallest error in ppm) was retained when more than one annotation was available for a given mass feature. Because annotations are based on exact mass and, where available, molecular sum formula information, confidence corresponds primarily to Schymanski levels 4-5 (Schymanski et al., 2014).

### MeOH extraction of plant material for LC-MS analysis

Metabolites were extracted from 10 mg of freeze-dried plant material by adding 1 mL of 100% MeOH containing 0.8 mg of phenyl-β-D-glucopyranoside (Sigma-Aldrich), 40 ng each of isotopically labelled internal phytohormone standards-D4-SA (Santa Cruz Biotechnology, USA), 40 ng D6-JA (HPC Standards GmbH, Germany), 40 ng D6-ABA (Santa Cruz Biotechnology, USA), 8ng of D6-JA-Ile (HPC Standards GmbH, Germany), and 10 ng of trifluoromethyl cinnamic acid (ThermoFisher Scientific Darmstadt, Germany CAS: 16642-92-5). Samples were shaken for 30s in a paint shaker (Skandex SO-10M, Fluid Management Europe, Netherlands) and then for 30 min at 200 rpm on a horizontal shaker (IKA Labortechnik, Staufen, Germany). After centrifugation, the supernatants were split for HPLC-UV and LC-MS/MS measurements and analysed subsequently. The different parameters (e.g. dilution, column, solvent, and gradient detector settings) for the different compound groups (salicinoids, phytohormones and phenolic acids, amino acids, free sugars) are described in the Supplemental Methods section.

### Analysis of isoprene synthase activity

Isoprene synthase (ISPS) activity was determined from 50 mg ground leaf powder (Bertić et al., 2023). Samples were mixed with Polyclar, extraction buffer (Tris/HCl with MgCl_2_, CaCl_2_, glycerol, PEG, and Tween 20; pH 8.0), and reducing agent, then incubated on ice with repeated vortexing. Extracts were centrifuged (18,000 rpm, 20 min). Desalting was performed using NAP-5 columns equilibrated with ISPS buffer (Tris/HCl with MgCl_2_ and glycerol, freshly supplemented with DTT). Enzyme reactions were initiated by adding DMADP and incubated for 1 h at 40 °C. Headspace isoprene was quantified immediately after incubation (details in Supporting Methods).

### RNA sequencing and transcriptome analysis

RNA extraction was performed as described by Geraldes et al. (2011), with subsequent RNASeq library construction carried out according to the Illumina RNA-Seq protocol (Illumina, Inc. San Diego, CA). RNA-Seq Libraries were sequenced on the Illumina HiSeq 1500 Sequencing System at Canada’s Michael Smith Genome Sciences Centre (Vancouver, BC) as 2 × 150 bp paired-end reads to a depth of ∼20-40 M read pairs per library. Demultiplexed FASTQs were quality-checked using FastQC for per-base quality, adapter contamination, and duplication metrics and were adapter- and quality-trimmed before alignment. Reads were mapped to the v3.0 *Populus trichocarpa* reference genome (Nisqually-1). Gene expression data are available in NCBI GEO under accession XXXXXX (ID to be added prior to final publication) using TopHat and normalised sequence abundance was calculated using Cufflinks, as described by Hefer et al. (2015).

### Calculated climate parameters from ClimateNA

Climate data for each provenance’s place of origins were obtained with ClimateNA (ClimateNA v7.60, Centre for Forest Conservation Genetics, Canada) for the 1981–2010 normal period (Wang et al., 2016). ClimateNA provides locally downscaled and spatially customisable climate data. We extracted mean annual temperature, mean temperature of the warmest and coldest month, annual and summer precipitation, and annual and summer heat-moisture index. Climate values represent site-level averages for the original location of each provenance. Pairwise Euclidean distance (d=i=1∑n(ai−bi)²) was calculated subsequently.

## Statistical analysis

For descriptive analyses, the Pearson correlation coefficient and the related *p*-value were calculated; R^2^ is the squared Pearson correlation coefficient.

Data were transformed logarithmically (log10), centred, and then scaled (Pareto) before multivariate analysis (Eriksson, 2013; Van Den Berg et al., 2006).

Discriminant Analysis of Principal Components (DAPC) was calculated using the R package ’adegenet’ (Jombart, 2015). Heatmaps were calculated using the heatmap() function from base R.

Orthogonal partial least squares discriminant analysis (O2PLS-DA) for the metabolomic or RNA-seq data was calculated using SIMCA P (Version 13, Sartorius SG, Göttingen, Germany). The discriminant model was computed with origin as the assigned discriminating variable codex.

Partial Least Squares (PLS) with metabolomic data was also calculated using SIMCA P (Version 13). The discriminant model was computed with an assigned discriminating variable codex corresponding either to origin, to the extreme-latitude origins (3 most northern and 2 most southern) genotypes, or to origins with latitudes above 51° N or below 49° N. Discriminant mass features were defined as loadings of the PLS model with a VIP value > 1. The discriminant mass features of the model with extreme-latitude origins were used to generate a PLS model for origins with latitudes above 51° N or below 49° N.

To compare climate conditions among origins, Euclidean distance between origins was calculated using the yearly weather parameters exported from ClimateNA. All temperature-dependent parameters were scaled (e.g. mean temperature, extreme temperature, etc.) before pairwise Euclidean distance was calculated for each combination, resulting in 231 values. Pairwise geographic distance between origins was estimated using Google Earth (Version 7.3, 2026).

We then calculated the pairwise Euclidean distance/Bray-Curtis distance for the different mass features. To do so, we averaged each mass feature for each origin and then scaled the data (Pareto), which also resulted in 231 values. Thus, for every pair-wise comparison, we obtained Euclidean distance for the mass features, climate parameters, and geographic distance. Correlation coefficients (and *p*-values) among these three parameters were then calculated using the Mantel test.

## Results

### Geographic origin shapes the *Populus trichocarpa* leaf metabolome

We used twenty-two different *P. trichocarpa* provenances originating from locations spanning over 1,200 km along the west coast of North America within the native range of the species. Most provenances originated in British Columbia, Canada, but two were from Oregon, USA. Many provenances were collected from riverbanks, and in some cases more than one provenance was sampled along the same river. For instance, the HOMB, HOMC, and HOMD provenances all originated along the Homathko River. Clonal replicates (cuttings) were used to establish a common garden at the University of British Columbia in Vancouver, BC (Fig 1d).

Four individual *P. trichocarpa* genotypes from each provenance were sampled in the common garden for metabolomic profiling of leaves and subsequent O2PLS-DA analysis (Fig. 1a). For this supervised ordination, we used 1,030 mass features that were consistently detected by FT-ICR-MS. The first two O2PLS-DA dimensions indicated grouping of the genotypes by place of origin (colour-coded in Fig. 1), suggesting distinct metabolomes related to geographic origin or genetic differences.

This pattern became more apparent when the analysis was performed using only 41 genotypes from 16 provenances (Fig. 1b), for which RNA-seq data were also available. Provenance groups tended to be closer to each other in the O2PLS-DA model as the geographic distance between them decreased, and provenances that were geographically close were assigned similar colours in Fig. 1. For instance, the green colours representing the southernmost geographical origins (PITS, FNYI, and HRSO) are located towards the top of the second dimension (y-axis) in Fig. 1b.

### RNA-seq profiles show extensive overlap among provenances in supervised ordination

RNA-seq profiles showed extensive overlap among provenances in supervised ordination (Fig 1c). Using the RNA-seq dataset (comprising 41 genotypes), we examined whether provenances could be differentiated based on transcriptome-wide expression patterns when analysed using the same supervised framework. Unlike the metabolomic profiles, O2PLS-DA of the RNA-seq data revealed little separation among provenances (Fig. 1c). In accordance with negative Q_(cum) values, the model based on transcriptome data was unable to predict provenance classification. Consequently, we did not consider RNAseq data towards analyzing transcriptomic separation by origin. Instead, we used RNA-seq to examine pathway-internal coherence and candidate biosynthetic gene modules.

### Annotated compound classes highlight flavonoids and isoprenoids

Metabolites annotated through the MONA database as flavonoids or isoprenoids were aggregated by summing feature intensities belonging to each class for each individual leaf, providing an estimate of total flavonoid and isoprenoid content. In the subset of 41 genotypes for which both metabolomic and transcriptomic data were available, we observed a clear and highly significant positive correlation between total flavonoid and total isoprenoid pools (Pearson R^2^ = 0.827, p < 0.01; Fig. 2A), indicating that genotypes with larger isoprenoid pools also tended to accumulate larger flavonoid pools.

**Fig 2.**
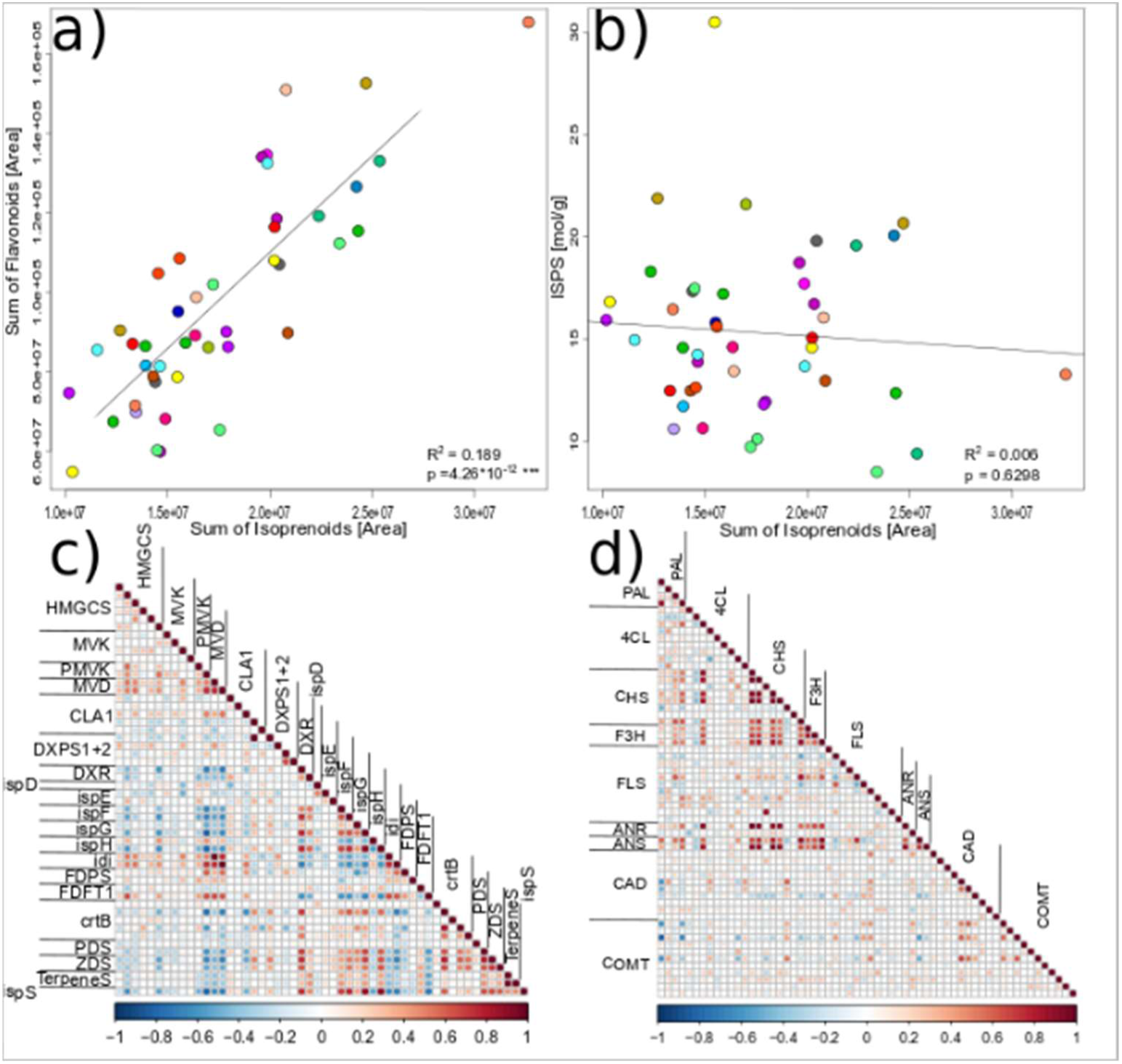
Correlation analysis comparing isoprenoid abundance, flavonoid abundance, isoprene synthase activity, and transcript abundance of putative biosynthetic genes. a) Correlation plot comparing the sum of all isoprenoids [summed feature areas on the x-axis] with the sum of all flavonoids [summed feature areas on the y-axis]. A linear regression line has been added (R^2^=0.189, p < 0.0001); b) Correlation plot of the sum of all isoprenoids [summed feature area] with isoprene synthase activity (ISPS) [µkat/kg]. Linear regression suggests no correlation (R^2^= 0.006, p = 0.6298). c) and d) Correlations matrix for selected genes putatively coding for biosynthetic steps in the terpenoid/isoprenoid c) and flavonoid d) pathways. Pearson correlation coefficients for each pairwise comparison is colour-coded as indicated underneath the matrix.

Limitations associated with very weak annotation certainties (level 5), precluded us from analysing the data further to identify the main drivers of this correlation at the level of individual compounds.

### Isoprene synthase activity does not track geography, climate, or total isoprenoids

Isoprene is the simplest isoprenoid and is synthesised from the same precursor, dimethylallyl pyrophosphate (DMAPP), as all other isoprenoids. The branch-point enzyme isoprene synthase (ISPS) converts DMAPP into isoprene (plus pyrophosphate), which could potentially divert carbon away from broader terpenoid metabolism.

As a proxy for isoprene emission capacity at the time of sampling, we quantified enzymatic ISPS activity in the same leaves that were used for non-volatile metabolomic profiling, because it was not feasible to measure emission directly for individual trees grown under field conditions. Although ISPS activity varied by more than two-fold across individuals, it did not exhibit any consistent climatic or geographic pattern across provenances. Moreover, despite an almost four-fold variation in total isoprenoid pools, ISPS activity did not predict overall isoprenoid accumulation, as there was no significant correlation between ISPS activity and total isoprenoid content (Pearson R^2^ = -0.07, p = 0.629; Fig. 2b). Together, these results suggest that variation in accumulated isoprenoids cannot be explained by competition with isoprene biosynthesis, and that ISPS activity itself does not exhibit a detectable climate- or distance-related trend within our samples.

### Chemodiversity differs among origins

Chemodiversity was quantified using the Functional Hill Diversity metric, which integrates richness, evenness, and chemical similarity within a given sample (Petrén et al., 2023). Functional Hill Diversity varied among provenances (Supplementary Fig. S2), suggesting that origins differ not only in class-level pools but also in how chemical space is structured within leaves.

From the transcriptome dataset, we selected genes with major roles in isoprenoid (Fig. 2c) or flavonoid (Fig. 2d) biosynthesis. We then compared transcript abundance within each pathway using pairwise Pearson correlation analyses. Despite the lack of correlation between ISPS activity and isoprenoid content, we observed positive correlations between expression of the *ispS* gene and that of two monoterpene synthases (labelled ’TerpeneS’ in Fig. 2c), as well as with *dxr* and *ispH* expression. The *dxr* gene encodes 1-deoxy-D-xylulose 5-phosphate reductoisomerase and is positioned close to the entry point into the MEP pathway. The *ispH* gene (also known as HDR) encodes 4-hydroxy-3-methyl-but-2-enyl pyrophosphate reductase and catalyses the final step in the pathway. Both enzymes play key regulatory roles in the MEP pathway. More generally, transcript abundance of genes in the two isoprenoid precursor pathways, namely the plastidial MEP pathway and the cytosolic pathway contributing to DMAPP, was positively correlated within each pathway, but negatively correlated between pathways (see Fig. 2c).

Within the flavonoid pathway, we observed strong positive correlations in transcript abundance between *PAL*, *CHS*, and *F3H*, as well as between *ANR* and *ANS*. *PAL* encodes phenylalanine ammonia lyase and catalyses the entry point into the general phenylpropanoid pathway, whereas *CHS* (chalcone synthase) catalyses the entry point into the flavonoid-specific branch. The other two enzymes also play important roles as branch-point enzymes in the biosynthesis of diverse classes of flavonoids. Only a comparatively small number of genes involved in other phenylpropanoid branch pathways were identified as co-expressed, which may reflect limited activity of these pathways in leaf tissue, for example towards lignin biosynthesis. Consistently, genes involved in the monolignol biosynthetic pathway that are present in our collection (e.g. *CAD* (cinnamyl alcohol dehydrogenase) or *COMT* (caffeic acid O-methyltransferase) were not detectably co-expressed with genes specific to the flavonoid pathway, although individual *CAD* and *COMT* genes were co-expressed with each other. Genes specific to other abundant soluble phenolics found in leaves, such as salicinoids or hydroxycinnamoyl conjugates, were not included in our gene selection.

When gene expressions were compared across pathways, no obvious correlations were observed between flavonoid- and isoprenoid-related genes. Likewise, no clear correlation was observed when total isoprenoid or total flavonoid abundance was compared with transcript abundance of the associated biosynthetic genes.

### Phenolic compounds show individual differences but no consistent provenance gradient

Abundance analysis of selected phenolics suggested river-system-related shifts in the targeted metabolite profiles. Visual differences were most apparent for benzoic acid (BA) and salicinoid-related compounds, whereas salicylic acid (SA) and the hydroxycinnamic acids showed more moderate shifts among origins (Fig. 3). In contrast, gallic acid and procyanidin B1 (PAB1) varied over a comparatively narrower range. Exploratory Kruskal-Wallis tests indicated nominal differences among river systems for some compounds, particularly ferulic acid, SA, gallic acid, and *p*-coumaric acid (p = 0.016-0.031), but these effects did not remain significant after correction for multiple testing. We therefore interpret these differences primarily as suggestive evidence for river-system-related variation. Abundance variation patterns for the remaining compounds are shown in the supplement (Sup. Fig. S3-S5).

**Fig 3:**
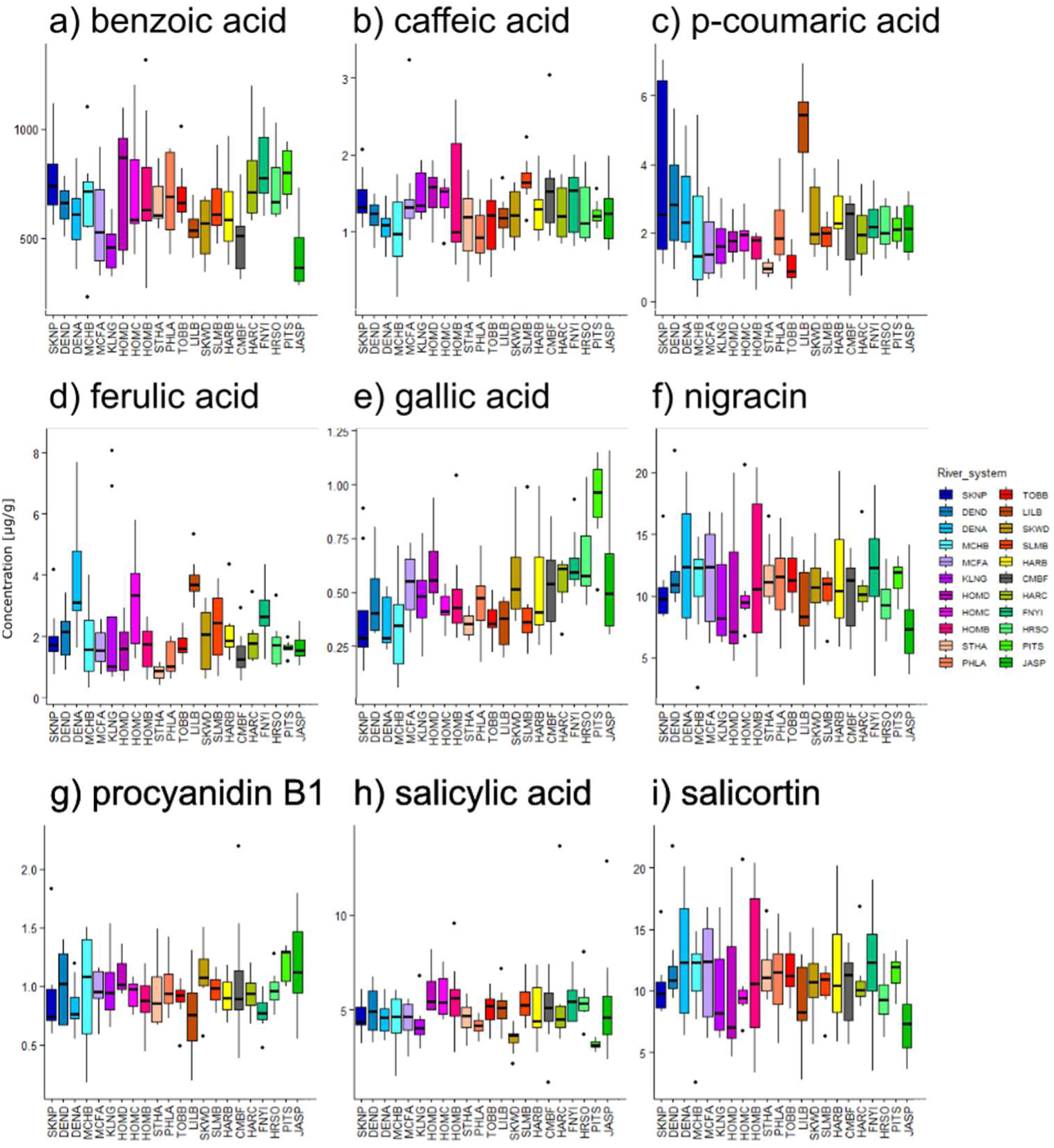
Selected targeted compounds per provenance. Boxplots for selected targeted compounds over provenances: a) Benzoic Acid (BA), b) Caffeic Acid, c) *p*-Coumaric Acid, d) Ferulic Acid, e) Gallic Acid, f) Nigracin, g) PAB1, h) Salicylic Acid, and i) Salicortin. All showing differences between provenances but no trend can be observed.

### Climate is an important factor explaining metabolomic differences

To determine whether geographic distance and climatic differences at the places of origin affect metabolic diversity, we analysed every possible pair of genotypes, calculating geographic distance in kilometers, Euclidean climatic distance based on multiple climate variables (temperature, precipitation and heat-moisture), and metabolomic dissimilarity using Bray-Curtis analyses. In each case, smaller distance measures indicate greater similarity between the genotypes compared. When Bray-Curtis dissimilarity was compared with either Euclidean climatic distance (Fig. 4a) or geographic distance (Fig. 4b), a weak but highly significant positive correlation became apparent. These correlations became more pronounced when the analyses were restricted to the southernmost and northernmost genotype origins only (Figs 4c and 4d), suggesting that latitudinal extremes play a disproportionately important role in the chemical differentiation within this collection. Overall, genotypes originating from locations that are more distant, either geographically or climatically, tend to exhibit greater differences in metabolomic composition than genotypes originating from geographically close regions with more similar historical climates. This pattern cannot be explained by acclimation, because all genotypes were grown in a common garden for several years before sampling. Collectively, these analyses support distance-decay of metabolomic similarity with both climate and geography of origin.

**Fig 4:**
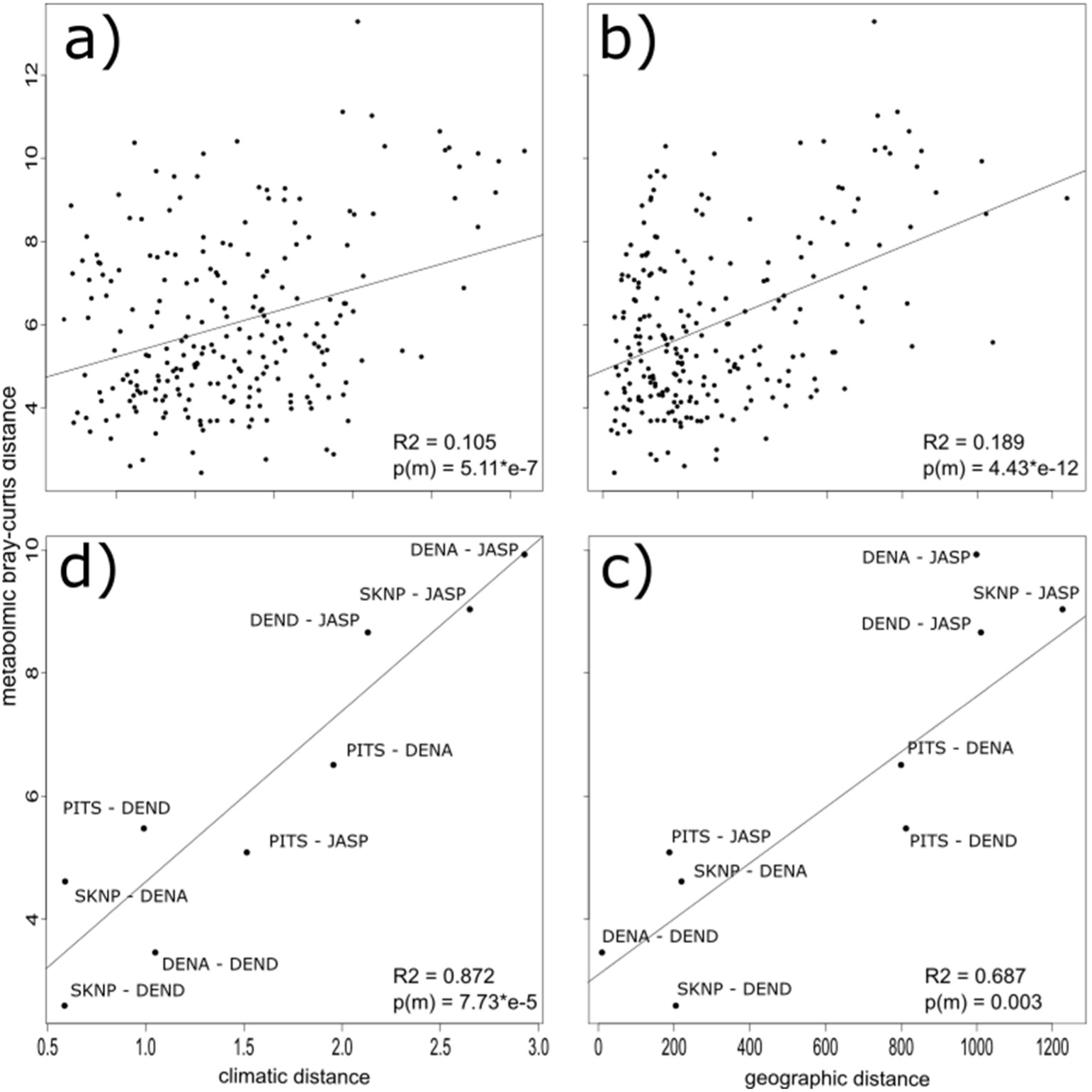
Metabolomic β-diversity in leaves of *P. trichocarpa* correlations with climate and geographic distances. Correlation between the Bray-Curtis distance of the metabolomic data of genotype pairs and the a) Euclidian climatic distance (R^2^=0.105, p(m) < 0.001) or b) geographic distance (in km) of the same pair. A linear regression line was added to each scatter plot. The lower plots show correlation analysis based on only for the two most southern (JASP, PITS) and three most northern (SKNP, DENA, DEND) provenances, between the Bray-Curtis distance of the metabolomic data and c) the Euclidian climatic distance (R^2^=0.872, p(m) < 0.001) or d) the geographic distance (in km) (R^2^=0.687, p(m) = 0.003).

### Metabolic diversity is governed by geographic distance of places of origin, even when genotypes are grown in a common garden

To further dissect differences between the southernmost and northernmost provenances, we employed Discriminant Analysis of Principal Components (DAPC) for the two southernmost (JASP and PITS) and three northernmost (SKNP, DENA, and DEND) provenances based on metabolomic data (Figure 5). The northern and southern provenance groups clearly separate along the x-axis (Fig 5a; circles represent 90% confidence intervals). Along the y-axis, the southern provenances are further separated, whereas the northern provenances separate less strongly, especially DENA and DEND, which both originate from the Dean River drainage basin. Figure 5b shows analogous separation of the same provenances based on RNAseq data and DAPC (excluding PITS, for which no RNA-seq data was available).

**Fig 5.**
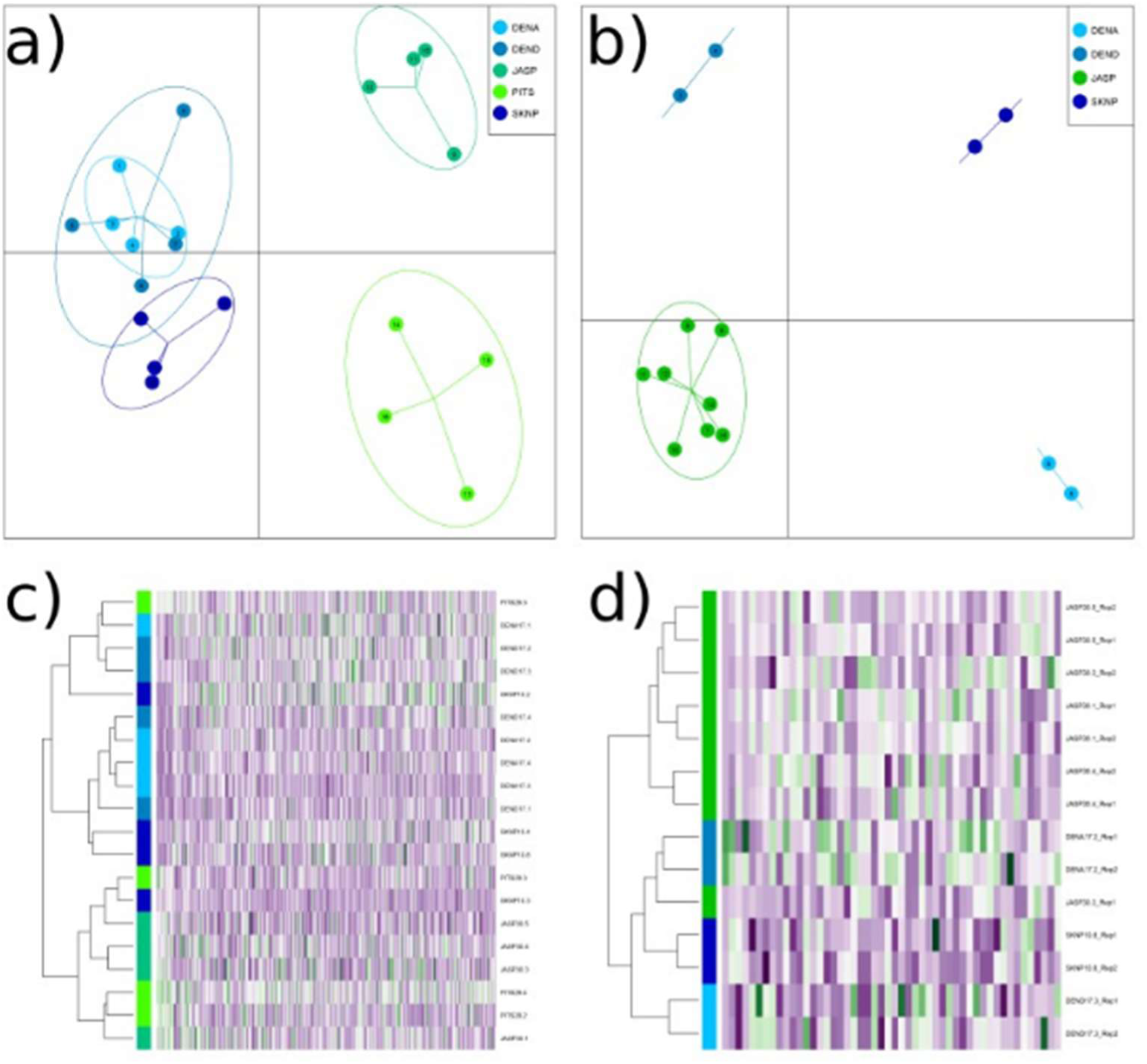
Discriminant Analysis of Principal Components (DAPC) of XY using most southern and northern provenances of *Populus trichocarpa*. a) DAPC of the two most southern (JASP, PITS) and three most northern (SKNP, DENA, DEND) provenances using the metabolomic data. Circles representing 90% confident intervals. Along the x-axis the southern and northern origins are well separated, while along the y-axis provenances are further separated except for DENA and DEND, which originated from the from the same river drainage (Dean River). b) DAPC based on RNAseq data using the same provenances (except PITS for which no RNAseq data were available). c) Separation of the metabolomic data using hierarchical clustering and visualisation as a heatmap (selected according to VIP-values > 1). d) Hierarchical clustering using the RNAseq data as described for the metabolomic data. The southern group (seven clonal replicate trees of the JASP provenance coloured green) separates well from the northern provenances.

Complementary hierarchical clustering of features selected according to VIP values >1 also shows that northern provenances (green hues) are generally well separated from southern provenances (blue hues) (Fig 4c). However, one genotype from each group clusters in the opposite group, and individual provenances within each group are not well resolved. Hierarchical clustering of the RNA-seq data likewise shows that the southern group (represented by seven clonal replicate trees of the JASP provenance; coloured green) is clearly distinct from the northern provenances (represented by two clonal replicates each for three provenances).

Separation of northern and southern provenances based on metabolite composition was further supported by PLS analysis, initially using the same subset of origins as in the DAPC and restricting the model to mass features with VIP scores >1 (Supplementary Material Fig. S4). Expanding the PLS model to include all provenances while retaining these features maintained a clear separation between northern and southern groups (Supplementary Material Fig. S5). In this broader analysis, the two Oregon State provenances (PITS and JASP) are further distinguished from other southern lines along the second component. Notably, these two provenances originate from latitudes below 46° N, whereas the remaining southern provenances originate above 49° N.

We then applied hierarchical clustering to the same set of discriminant mass features (VIP >1) and visualised the results in a heatmap alongside RNA-seq data for the northernmost and southernmost provenances (JASP, PITS, SKNP, DENA, and DEND). Based on metabolomic profiles, samples segregate strongly by geographic origin, with northern (SKNP, DENA, DEND) and southern (JASP, PITS) provenances forming well-defined clusters. Only two individuals (PITS 29.4 and SKNP 10.3) group with the opposite cluster. In contrast, RNA-seq data show weaker separation among provenances and less consistent clustering by geographic origin. The two Dean River provenances tend to cluster in proximity in metabolomic data, although this pattern is less pronounced than in the DAPC and PLS analyses.

### River-drainage structure within regions

Next, we selected groups of provenances originating from the same river drainage to further test whether geographical connectedness defines metabolic similarity. The three river drainages compared were the Dean River (DENA, DEND) in the north, the Homathko River (HOMA, HOMB, HOMD) in the north-central region, and the Harrison River (HARB, HARC) in the southern region of British Columbia. These drainages are at least 200 km apart, whereas provenances within a given drainage originated no more than 40 km apart. DAPC analysis revealed clear separation by river along the first discriminant axis (see Fig. 6a-b), although the Dean and Homathko groups overlapped in the metabolomic DAPC analysis (see Fig. 6a). The two Harrison River provenances did not separate along the primary axis when using the metabolomic dataset but did so along the secondary axis. River systems were also clearly separated when focusing on the between-group variance of the transcriptome data in the DAPC analyses (Fig. 6b). As before, provenances from the same river system tended to be indistinguishable from each other. The exception was HOMC from the Homathko River, which was placed separately from HOMB and HOMD in the RNA-based DAPC. In contrast, clustering of highest contributing features or transcripts (VIP >1) did not consistently recover river groupings for either data type (Fig. 6c,d).

**Fig 6.**
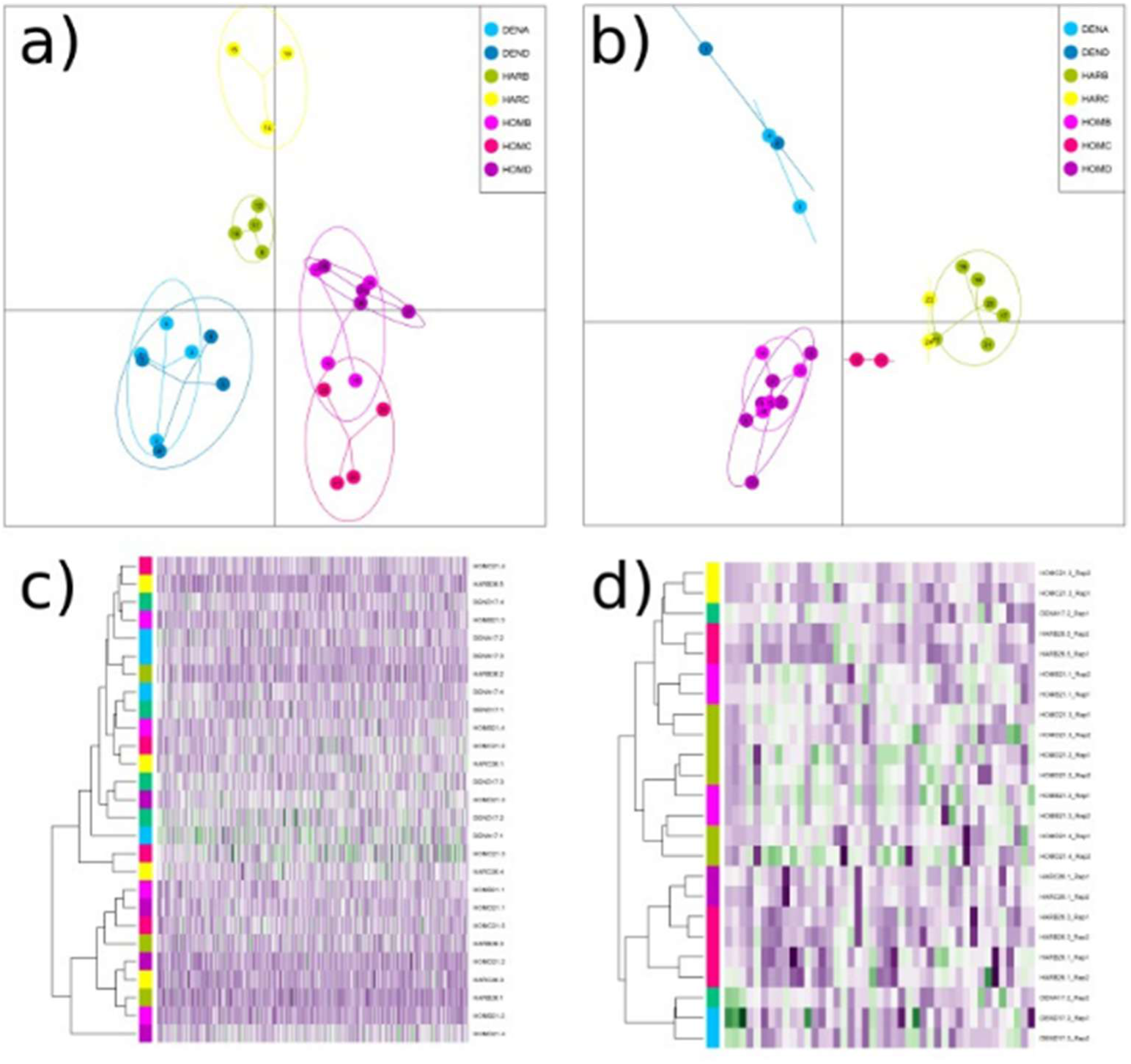
DAPC of Rivers and RNA Correlation. a) and b) DAPC of the three river systems (Homatoka, Dean, Harrison River) using the metabolomic data a) and the RNASeq data b), circles representing 90% confident interval; all the three river system can be distinguished. c) and d) Separation of the origins from a) in a heatmap using the metabolomic data c), (selected according to VIP-values > 1)) and the RNAseq data d), no grouping was apparent.

Taken together, both metabolic and transcriptomic diversity increase with geographic and climatic distance of origin, even when trees are cultivated in a common garden. Genotypes originating from nearby locations, such as those from the same provenance or river drainage, tend to be highly similar in their metabolic and transcriptomic composition, whereas signatures of diversity among provenances or river basins are readily detectable.

## Discussion

### Coordinated flavonoid-isoprenoid allocation points to shared stress-responsive regulatory control

The strong positive covariation between flavonoid and isoprenoid pools across *Populus trichocarpa* individuals grown under identical environmental conditions suggests that these pathways are not regulated independently but instead are coordinated through shared upstream regulatory processes. Because all genotypes were cultivated for years in a common garden, the observed covariance is unlikely to reflect short-term environmental acclimation alone and instead points to genetically encoded or developmentally stabilised allocation programs. Mechanistically, several lines of evidence support such coordination. Plastidial isoprenoid biosynthesis through the methylerythritol phosphate (MEP) pathway is tightly linked to cellular redox balance and abiotic stress signaling. Perturbation of MEP-pathway flux alters abscisic acid (ABA) biosynthesis and oxidative stress responses, both of which are known regulators of phenylpropanoid and flavonoid metabolism (Barta & Loreto, 2006; Monson et al., 2021; Businge et al., 2024). In poplar, RNAi-mediated suppression of isoprene emission reduces not only isoprenoid-derived products but also flavonoid accumulation, indicating that plastidial isoprenoid metabolism can influence phenylpropanoid allocation beyond simple precursor competition (Behnke et al., 2010). Conversely, ectopic isoprene production enhances accumulation of phenylpropanoids and antioxidant metabolites under stress conditions (Tattini et al., 2014). Together, these findings support a model in which plastidial stress sensing and redox signaling influence coordinated carbon allocation into both flavonoid and isoprenoid branches (Sugimoto et al., 2022; Cappellari et al., 2020; Monson et al., 2021).

Our data is consistent with this framework. Genotypes with larger isoprenoid pools also accumulated larger flavonoid pools, whereas isoprene synthase (ISPS) activity itself was unrelated to total isoprenoid abundance. This distinction is mechanistically important. If variation in accumulated isoprenoids were driven primarily by substrate competition between volatile isoprene formation and non-volatile terpenoid synthesis, a negative relationship between ISPS activity and total isoprenoid pools would be expected. Instead, the absence of such a relationship suggests that upstream regulatory nodes, rather than branch-point competition alone, determine coordinated metabolite allocation. The transcript data further supports this interpretation. Within the MEP pathway, expression of *dxr*, *ispH*, *ispS*, and terpene synthases showed positive covariance, consistent with coordinated regulation of plastidial isoprenoid biosynthesis. By contrast, genes associated with cytosolic DMAPP-contributing pathways were negatively correlated with plastidial MEP-module expression, suggesting regulatory partitioning between plastidial and cytosolic isoprenoid metabolism.

Such compartment-specific coordination has been proposed previously for poplar isoprenoid metabolism (Movahedi et al., 2022). Importantly, however, we detected little correspondence between transcript abundance and accumulated metabolite pools across pathways, indicating that metabolite accumulation likely integrates transcriptional, post-transcriptional, enzymatic, and substrate-level controls over longer timescales.

Collectively, these findings support a mechanistic model in which provenance-dependent differences in stress-responsive regulatory architecture influence carbon allocation into both flavonoid and isoprenoid metabolism. Rather than representing isolated pathway responses, the observed class-level covariance likely reflects integrated regulation mediated through plastidial stress sensing, ABA/redox signaling, and coordinated metabolic allocation.

### Transcript–metabolite decoupling reflects temporal and regulatory integration across biological levels

Although transcript abundances within biosynthetic pathways were strongly coordinated, transcriptomic patterns showed comparatively weak correspondence with accumulated metabolite pools. Such transcript–metabolite decoupling is common in specialised metabolism and reflects the fundamentally different temporal and regulatory properties of these biological layers.

RNA abundance captures a comparatively instantaneous cellular state, whereas many specialised metabolites accumulate over extended developmental periods and integrate biosynthetic activity over time. In woody perennials such as poplar, flavonoids, salicinoids, and many terpenoid-derived metabolites can remain stored within tissues long after transcript abundance has changed. Consequently, metabolite pools may reflect historical gene expression states rather than transcriptional activity at the time of sampling.

In addition, specialised metabolism is regulated at multiple mechanistic levels beyond transcription. Flux through the MEP and phenylpropanoid pathways depends on substrate availability, redox state, enzyme stability, compartmentalisation, transport processes, and post-translational regulation. Several enzymes in the MEP pathway, including DXR and ispH/HDR, are known regulatory nodes sensitive to oxidative and environmental conditions (Monson et al., 2021; Businge et al., 2024). Similarly, flavonoid biosynthesis is controlled through combinatorial transcription-factor networks and inducible signaling cascades involving jasmonate and ABA.

The within-pathway co-expression patterns detected here nevertheless indicate biologically coherent regulatory organisation. Positive correlations among core flavonoid genes (*PAL*, *CHS*, *F3H*, *ANS*, *ANR*) are consistent with coordinated regulation of flavonoid branch flux, whereas co-expression among MEP-pathway genes suggests integrated plastidial isoprenoid regulation. The lack of strong cross-pathway transcript coordination despite clear metabolite covariance therefore suggests that coordination emerges at higher-order regulatory or physiological levels rather than through simple synchronous transcription.

This interpretation has important implications for metabolomic studies in trees. Metabolite-based phenotypes may capture long-term integrated physiological strategies more effectively than snapshot transcript profiles. The stronger provenance structure in metabolomes compared with transcriptomes observed here supports this view and suggests that metabolomic organization may provide a particularly robust readout of inherited allocation strategies and stress-adaptive physiology (Redestig & Costa, 2011; Liang et al., 2016; Dubois et al., 2017).

### Provenance-dependent metabotypes persist under common-garden conditions

The persistence of a metabolomic imprint of geographic origin under common-garden conditions, the clear separation at latitudinal extremes, and the partial grouping by river drainage all suggest that factors linked to origin shape leaf chemical phenotypes. The observed decrease in similarity as geographic and climatic distance between sites of origin increases further supports this interpretation. As growth and sampling conditions were uniform, these differences most likely reflect heritable genetic variation and/or stable epigenetic states established at the site of origin prior to sampling. These states then persisted through clonal propagation from an earlier provenance trial. This interpretation is consistent with extensive evidence for local adaptation of plant phenotypes along climate gradients in common-garden settings (Eisenring et al., 2022), as well as with the general view that phenotypes arise from interactions between genotype and environment (G × E). In *P. trichocarpa*, multi-site common-garden and landscape-genomic studies have demonstrated that phenotypic and genetic structure track environmental gradients across the species’ range. Numerous loci have been implicated in ecological trait variation (McKown et al., 2014a, b; Porth et al., 2013b; Geraldes et al., 2014a, b). Furthermore, plastic responses across years within a common garden emphasize that temporally varying environments can modulate trait expression in addition to provenance effects (Liu & El-Kassaby, 2019). Similarly, physiological contrasts among allopatric Populus species planted in common gardens reveal geographic clines in photosynthesis, growth, and isotope discrimination consistent with climate-linked adaptation (Soolanayakanahally et al., 2015). Latitudinal transect studies in European aspen further contextualise north-south separations by linking clinal genomic variation to adaptive life-history differences and to a major phenology locus (*PtFT2*) (Ingvarsson & Bernhardsson, 2020; Wang et al., 2018). There is also evidence that adaptive introgression has facilitated high-latitude adaptation (Rendón-Anaya et al., 2021). Together, these findings reinforce the plausibility that the geographic metabolomic structure we recover reflects underlying allelic differences at loci influencing specialised metabolism and its regulation.

In contrast, we did not observe a consistent north-south gradient for the metabolic markers examined here, although patterns in the targeted metabolite dataset may indicate underlying population-level genetic or chemotypic differentiation in foliar metabolism, for example in salicinoids. Because all plants were grown under common-garden conditions, the observed variation is unlikely to reflect the immediate climatic environment and instead is more likely linked to inherited differences among source populations. Given the central role of these metabolites in poplar defence, the river-system-specific patterns we observe may therefore reflect differences in constitutive defence-related metabolism rather than immediate environmental effects (Ullah et al., 2019).

Beyond *Populus*, an increasing body of work on forest adaptation shows that provenance and climate of place of origin consistently influence performance across sites, and that these factors can inform climate-smart deployment. Reviews and syntheses of North American provenance and common-garden trials reveal robust origin effects that can be generalised, providing templates for assisted migration policies (Park & Rodgers, 2023; Stanturf et al., 2024; Bower et al., 2024). Multi-species analyses indicate that growth stability under novel climates is predicted by the climate at the origin of the provenance (Di Fabio et al., 2024), which aligns with our finding that metabolomic similarity decreases with climatic and geographic distance of origin. Modelling suggests that assisted migration could help preserve the forest carbon sink in the face of climate change (Chakraborty et al., 2024), while ’genomic offset’ frameworks integrate genotype-climate associations with niche models to forecast population vulnerability (Chen et al., 2022). However, recent benchmarking cautions that genomic predictors vary in performance and should be validated against reciprocal transplant or long-term common-garden datasets (Lind et al., 2024). In this context, our provenance-linked metabolomic structure offers a complementary phenotype that could augment trait- and genomic-based approaches for guiding seed transfer in a changing environment. Biogeographic projections of floristic reorganisation also highlight the importance of conserving provenance-linked functional and chemical diversity (Minev-Benzecry et al., 2024).

### Functional chemodiversity reflects organisation of chemical space rather than abundance of individual compounds

A substantial body of research on *Populus* species beyond *P. trichocarpa*, including studies on trembling aspen (*P. tremuloides*) and European aspen (*P. tremula*), shows that defence chemistry is influenced by genotype and environment, and that the ecological impact of chemical variation can be substantial. Early experiments revealed that the interaction between genotype, nutrient availability, and defoliation determines aspen phytochemistry and herbivore performance. More broadly, the efficacy of chemical defence depends on genetics, environment, and their interaction (G × E) (Osier & Lindroth, 2001; Donaldson & Lindroth, 2007). Reviews have emphasised the central roles of salicinoids and condensed tannins, while also highlighting inducibility, costs, and context dependence (Lindroth & St. Clair, 2013). More recent work has linked foliar trait syndromes to explicit growth-defence trade-offs, demonstrating that constitutive allocation rules can lead to divergent growth trajectories even in the absence of significant herbivory (Kruger et al., 2020). However, not every trait manipulation propagates to community-level differences. For instance, altering lignin in hybrid poplar did not substantially modify plant defence or arthropod communities in one study (Buhl et al., 2017). We therefore adopt a measured stance: genotype and environment jointly structure chemistry and its ecological effects, but the magnitude and direction of these effects vary across species, compound classes, spatial scales, and ecological contexts. Within this broader perspective, our finding that flavonoids and isoprenoids co-vary while metabolomes separate by provenance under shared growth conditions aligns with a general pattern of coordinated investment and heritable chemical setpoints.

### Genetic architecture, chemical organisation, and why chemodiversity matters

Genetic mapping in *Populus* has revealed loci with large effects on key defence compounds, such as salicinoids in hybrid cottonwoods. This implies that differences in allele frequency at a limited number of loci can substantially reshape leaf chemical space (Woolbright et al., 2018). Such architecture provides a plausible basis for the geographic and climatic differences observed here: landscape sorting of alleles at defence-relevant loci would be expected to generate latitudinal and drainage-level metabolomic differentiation that persists in a common environment. However, single-compound markers are often labile or pleiotropic, and may not reflect broader geographic patterns. For example, salicinoid chemotypes show limited geographic structure in parts of the *P. tremula* range (Keefover-Ring et al., 2014). These contrasts strengthen the case for focusing on multivariate chemical organisation.

Functional Hill Diversity, which considers richness, evenness, and biochemical/structural similarity, differed among origins in our data and helps explain why targeted subsets of compounds produced weaker geographic signals than the untargeted feature space (Petrén et al., 2023). Theory linking phytochemical diversity to herbivore deterrence, enemy confusion, and robustness to multi-stressor environments, together with experimental evidence that higher chemical diversity can enhance defence against herbivory, suggests that chemodiversity is a functional trait with ecological relevance (Wetzel & Whitehead, 2020; López-Goldar et al., 2024). Recent conceptual work has also started to formalise how chemodiversity should be quantified and modelled across ecological and evolutionary contexts (Hanusch et al., 2026 preprint), thereby strengthening the theoretical basis for treating chemodiversity as an interpretable trait, rather than a purely descriptive index (Thon et al., 2024). GWAS studies of poplar metabolites that implicate family 1 UDP-glycosyltransferases in specialised metabolite variation further support the view that genetically encoded differences in late-step tailoring enzymes can reorganise chemical space and, by extension, influence functional chemodiversity (Saint-Vincent et al., 2023).

## Methodological considerations

The untargeted FT-ICR-MS approach provides ultrahigh mass resolution and broad coverage, but it predominantly yields exact-mass and formula-level information. Consequently, many features remain at Schymanski identification levels 4-5 (Schymanski et al., 2014). Class-level aggregation and similarity-based chemodiversity metrics should therefore be interpreted as conservative descriptors of feature space rather than definitive structural inventories. Greater use of MS/MS confirmation and pathway-informed annotation will improve mechanistic interpretation and facilitate the identification of marker compounds. Taken together, our results indicate that *P. trichocarpa* leaf chemistry encodes both short-term regulatory programs linked to stress physiology and longer-term evolutionary and ecological histories tied to provenance. The coordinated variation of flavonoid and isoprenoid pools, pathway-internal transcript coherence with cross-pathway decoupling, the imprint of geographic and climatic origin, and among-origin differences in Functional Hill Diversity collectively support an interpretation in which heritable allocation rules and stress-linked regulation shape metabotypes in ways that are likely to matter for resilience in the face of environmental change.

## Further directions

Longitudinal sampling to quantify ISPS activity, isoprene emission, class-resolved metabolite pools, chemodiversity metrics, transcript and protein states, and growth responses to controlled challenges and natural variation would allow causal inference about links among MEP-mediated stress sensing, ABA/redox signaling, phenylpropanoid allocation, and the growth-defence balance described by the isoprene-centred theory (Monson et al., 2021). In parallel, placing multiple provenances across replicated common gardens along climatic gradients, complemented by genomic analyses to identify defence-related loci, would facilitate the conversion of metabolomic structure into actionable guidance for provenance selection and assisted gene flow in the context of climate change (Eisenring et al., 2022; McKown et al., 2014a, b; Porth et al., 2013b; Liu & El-Kassaby, 2019; Soolanayakanahally et al., 2015).

Assisted migration frameworks offer practical ways to turn the results of such trials into management strategies (Park & Rodgers, 2023; Stanturf et al., 2024; Bower et al., 2024). Recent analyses suggest that these methods could help to preserve forest carbon reserves in the face of climate change (Chakraborty et al., 2024). However, given the variable predictive performance of genomic offset metrics (Lind et al., 2024), metabolomic fingerprints, particularly class-level compound pools and Functional Hill Diversity, offer a complementary heritable readout that can be evaluated against survival, productivity, and resilience across environments. Furthermore, projections that climate change will reorganise and potentially homogenise global floristic regions (Minev-Benzecry et al., 2024) emphasise the importance of conserving provenance-linked chemical variation in order to sustain functional diversity and ecological interactions. Therefore, determining the extent to which chemodiversity reflects adaptive potential represents a key next step (Chen et al., 2022; Loreto, 2024).

Overall, our results suggest that the leaf chemistry of *P. trichocarpa* encodes short-term regulatory programs associated with stress physiology, as well as longer-term evolutionary and ecological histories linked to provenance. The coordinated variation of flavonoid and isoprenoid pools, coherence of transcripts within pathways with decoupling across pathways, the imprint of geographic and climatic origin, and differences in Functional Hill Diversity among origins collectively support the idea that heritable allocation rules and stress-linked regulation shape metabotypes in ways that will be important for resilience in the face of ongoing environmental change.

## Acknowledgements

This work is dedicated to the memory of Carl J. Douglas, UBC, Vancouver, who was a friend, mentor, and an outstanding scientist. Without him, this work would never have been created. This work was supported in part by the Genome Canada Large-Scale Applied Research Project (168BIO) awarded to JE and SDM. Additional support was provided by the DFG through Research Unit FOR3000 to JPS and SBU (SCHN563-8/1-2; UN276-5/1-2).

During the preparation of this work, the author(s) used AI tools (such as ChatGPT 5.4 and DeepL 1.85.0) in order to polish the writing framework and search relevant research articles (literature overview). After using this tool/service, the author(s) reviewed and edited the content as needed and take(s) full responsibility for the content of the published article.

## Author contributions

JPS, SDM, JE, and SU designed the study. AM, SDM, and JE collected the plant material. IZ performed sample preparation and enzymatic analyses. CH and JE conducted transcriptomic analyses. MP, SJV, BK, and PSK performed metabolomic analyses and statistical analyses. MP wrote the manuscript with input from SJV, JE, SU, and JPS. All authors reviewed, revised, and approved the final manuscript.

## Data availability

The transcriptomic raw data will be available in the NCBI GEO under accession XXXXXX (ID to be added prior to final publication). All processed datasets, including metabolite abundance matrices, and metadata required to reproduce the analysis, are archived on Zenodo https://doi.org/10.5281/zenodo.20049138.

## Supporting information

Supporting Material and Methods

Figure S1: Van Krevelen-diagram coloured according to MONA-Annotation

Figure S2: Functional-Hill Diversity between origins

Figure S3: Additional phythormones between origins

Figure S4: Sugar compounds between origins

Figure S5: Amino acids between origins

Figure S6: DAPC for targeted compounds for the North-South extremes according to Fig. 5

Figure S7: DAPC for targeted compounds for the North-South extremes according to Fig. 6.

Table S1: Details of analysis of phytohormones by LC-MS/MS (HPLC 1260 (Agilent Technologies)-QTRAP6500 (SCIEX)) in negative ionisation mode.

Table S2: Details of analysis of amino acids by LC-MS/MS (HPLC 1260 (Agilent Technologies)-QTRAP6500 (SCIEX)) in positive ionisation mode.

Table S3: Annotations of the MONA-Database - external provided in .xlsx-File.

