## Supplementary for "Geographic and Climatic Origins Shape the Leaf Metabolome of *Populus trichocarpa*"

### Supplemental Material

### Supplemental Methods

*Salicinoid analysis*

To 400 µL of the raw extracts (see above) 400 µL of Milli-Q H2O was added before measuring the analytes using high-performance liquid chromatography-ultraviolet detection (HPLC-UV, Agilent 1100 HPLC system, Agilent Technologies).Then, 10 µL of the diluted extract was injected into a reversed-phase chromatographic column (EC 250 × 4.6 mm NUCLEODUR Sphinx RP, 5 µm, Macherey Nagel, Germany) connected to a pre-column (C18, 5 µm, 4×3 mm, Phenomenex, USA). The mobile phases consisting of two solvents, solvent A (Milli-Q H2O) and solvent B (acetonitrile), were run in gradient mode. The time/concentration (min/%) of the gradient was set to 0/14; 22.00/58; 22.10/100; 25.00/100; 25.10/14; 30.00/14 with a constant flow rate of 1 mL $min^{-1}$. The column oven temperature was set to 25 °C. The signal at 200nm and 280nm was detected with a photo diode array detector (PDA, Agilent 1100 DAD). Using these settings, the retention times (RT) of the compounds of interest were 5.2 min (salicin), 6.4 min (PAB1), 10.4 min (salicortin), 12.6 min (nigracin), 13.25 (trichocarpin), and 17.91 (tremulicin). Data acquisition and processing was accomplished using DataTrans.exe software.

*Phytohormone, and phenolic acids analysis*

Phytohormones and phenolic acids were analyzed via HPLC (Agilent 1260 series HPLC system (Agilent Technologies)) coupled to a triple quadrupole mass spectrometer (QTRAP6500 (SCIEX, Darmstadt, Germany)). The analytes were injected onto a UPLC column (Zorbax Eclipse XDB-C18 column, 1.8 µm, 4.6×50 mm, Agilent Technologies, USA) connected to a pre-column (C18, 5 µm, 4 × 3 mm, Phenomenex). The injection volume was set to 2 µL. Two solvents, solvent A (0.05% formic acid in H2O) and solvent B (acetonitrile) were used. The following chromatographic gradient was applied: (time in min/concentration of solvent B in %): 0.00/5, 0.50/5, 9.50/58, 9.52/100, 11.00/100, 11.10/5 and 14.00/5. The constant flow rate was set to 1100 µL $min^{-1}$. The temperature of the column oven was set to 25 °C. Electrospray ionization (ESI) in negative ionization mode was used for the coupling of LC to MS. The mass spectrometer parameters were set as follows: ion spray voltage, -4500 V; turbo gas temperature, 700 °C; collision gas, 7 psi; curtain gas, 35 psi; ion source gas 1, 60 psi; ion source gas 2, 60 psi. Parent ion to product ion was monitored by multiple reaction monitoring (MRM) as listed in table S1. Data processing was performed using MultiQuant 3.0.3 software (SCIEX, Darmstadt, Germany) and analyte quantity was determined relative to the corresponding internal standard peak area. Concentration of cis-OPDA, and OH-JA were determined relative to the quantity of the internal standard D6-JA with a theoretical response factor one. OH-JA-Ile and COOH-JA-Ile were quantified relative to D6-JA-Ile with a theoretical response factor one. Sulfo-JA, OH-JA-Glucoside, and dinor-OPDA were determined relative to the quantity of the internal standard D6-JA with experimentally determined response factors of 6.0; 3.7; and 0.7, respectively.

*Amino acid analysis*

Free amino acids were analysed from the same raw extracts as used for phytohormone analysis. The raw extracts were diluted 1:10 with water containing an isotopically labelled amino acid mix (13C, 15N- labelled amino acid mix at a concentration of 10 µg of the mix per mL; from Isotec, Miamisburg, OH, USA). The extracts were measured with HPLC (Agilent 1260 series HPLC system (Agilent Technologies) coupled to a triplequadrupole mass spectrometer (QTRAP6500 (SCIEX, Darmstadt, Germany). Separation was achieved on a Zorbax Eclipse XDBC18 column (50 mm × 4.6 mm, 1.8 µm, Agilent Technologies, Germany). Two solvents, solvent A (MilliQ- Water) and solvent B (acetonitrile), were used. The following chromatographic gradient was applied: (time in min/concentration of solvent B in %): 0.00/3, 1.00/3, 2.70/100, 3.00/100, 3.10/3, 6.00/3. The constant flow rate was set to 1100 µL/min. The temperature of the column oven was set to 25 °C. Analytes were ionized in positive electrospray ionization mode. Multiple reactions monitoring (MRM) was used to monitor analyte parent ion → product ion: MRMs were chosen as in (Jander et al., 2004), see Suppl. Table S2, except for Arg (m/z 175 → 70) and Lys (m/z 147 → 84). Analyst 1.6 software (AB Sciex, Darmstadt, Germany) was used for data acquisition and MultiQuant 3.0.3 software (SCIEX, Darmstadt, Germany) was used for processing. Individual amino acids in the sample were quantified by the respective $^{13}C$, $^{15}N$-labelled amino acid internal standard.

*Analysis of free sugars*

For soluble sugar analysis the raw extracts of the samples were diluted 1:10 (v:v) in water. Glucose, fructose, sucrose and mannitol were measured on an Agilent 1200 series HPLC system (Agilent Technologies) coupled to an API 3200 mass spectrometer (AB SCIEX, Darmstadt, Germany) as described in Madsen et.al (2015). Separation was archived on apHera$^{TM}$ NH2 polymer HPLC column (15 cm × 4.6 mm, 5 µm; Supelco) by water (A) and acetonitrile (B) at a flow rate 1.0 ml/ min with an elution profile as follows: 0 to 0.5 min, 80% B; 0.5 to 13 min, 80% to 55% B in A; 13 to 14 min, 55% to 80% B; 14 to 18 min, 80% B. Electrospray ionization (ESI) in negative ionization mode was used for the coupling of LC to MS. The mass spectrometer parameters were set as follows: ion spray voltage, -4500 V; turbo gas temperature, 600 °C; collision gas, 5 psi; curtain gas, 20 psi; ion source gas 1, 50 psi; ion source gas 2, 60 psi. Precursor and product ions were monitored by scheduled MRM as following: m/z 178.8 → 89.0 (CE, -10 V; DP, -25 V) at 6.6 min for glucose; m/z 178.8 → 89 (CE, -7.5 V; DP, -30 V) at 5.6 min for fructose; m/z 340.9 → 59.0 (CE, -46 V; DP, -45 V) at 8.0 min for sucrose; m/z 180.9 → 89 at 7 min (CE, -22 V; DP, -35 V) for mannitol. Quantification of these carbohydrates was determined by external standard curves using commercial standards D-(-)-fructose, D-(+)-glucose and sucrose (all from Sigma-Aldrich).

### Supplementary Tables

**Supplemental Table S1.** Details of analysis of phytohormones by LC-MS/MS (HPLC 1260 (Agilent Technologies)-QTRAP6500 (SCIEX)) in negative ionisation mode.

| Q1 | Q3 | RT (min) | compound | Internal std | RF | DP | EP | CE | CXP |
|---|---|---|---|---|---|---|---|---|---|
| 136.93 | 93 | 5.9 | SA | $D_4$-SA | 1.0 | -20 | -8 | -24 | -7 |
| 263 | 153.2 | 6.4 | ABA | $D_6$-ABA | 1.0 | -20 | -12 | -22 | -2 |
| 209.07 | 59 | 7.2 | JA | $D_6$-JA+D5-JA | 1.0 | -20 | -9 | -24 | -2 |
| 140.93 | 97 | 5.9 | $D_4$-SA | | | -20 | -8 | -24 | -7 |
| 269 | 159.2 | 6.5 | $D_6$-ABA | | | -20 | -12 | -22 | -2 |
| 215 | 59 | 7.2 | $D_6$-JA | | | -20 | -9 | -24 | -2 |
| 214 | 59 | | $D_5$-JA | | | -20 | -9 | -24 | -2 |
| 121 | 121 | 5.6 | benzoic acid | TFMCA | | -20 | -10 | -5 | -10 |
| 163 | 118.9 | 4.7 | coumaric acid | TFMCA | | -20 | -8 | -20 | -5 |
| 179 | 134.9 | 4.0 | caffeic acid | TFMCA | | -20 | -8 | -22 | -5 |
| 193.1 | 133.9 | 5.0 | ferulic acid | TFMCA | | -20 | -8 | -22 | -5 |
| 169 | 125 | 1.3 | gallic-acid | TFMCA | | -20 | -10 | -18 | -19 |
| 215.06 | 171.05 | 7.3 | Trifluoro-methyl-cinnamic acid (TFMCA) | | | -20 | -8 | -18 | -4 |

**Supplemental Table S2.** Details of analysis of amino acids by LC-MS/MS (HPLC 1260 (Agilent Technologies)-QTRAP6500 (SCIEX)) in positive ionisation mode.

| Q1 | Q3 | RT (min) | Compound | DP | EP | CE | CXP |
|---|---|---|---|---|---|---|---|
| 90.1 | 44.1 | 0.5 | Ala | 51 | 5.5 | 17 | 4 |
| 106 | 60.1 | 0.5 | Ser | 61 | 4.5 | 15 | 4 |
| 116.1 | 70 | 0.5 | Pro | 61 | 7.5 | 19 | 4 |
| 118.1 | 72.2 | 0.5 | Val | 56 | 5 | 13 | 4 |
| 120.1 | 74.2 | 0.5 | Thr | 56 | 4.5 | 13 | 4 |
| 132.2 | 86.1 | 1.3 | Ile+Leu | 56 | 4.5 | 13 | 4 |
| 134.1 | 74.1 | 0.5 | Asp | 56 | 5.5 | 19 | 4 |
| 148.1 | 102.1 | 0.5 | Glu | 56 | 5.5 | 15 | 4 |
| 150.2 | 104.1 | 0.7 | Met | 51 | 4 | 13 | 4 |
| 156.2 | 110.1 | 0.4 | His | 61 | 5.5 | 17 | 4 |
| 166.2 | 120.2 | 2.6 | Phe | 56 | 6 | 17 | 4 |
| 175.1 | 70.1 | 0.4 | Arg | 66 | 6 | 31 | 4 |

| | | | | | | | |
|---|---|---|---|---|---|---|---|
| 182.1 | 136.2 | 1.4 | Tyr | 56 | 7 | 17 | 4 |
| 133.1 | 74.1 | 0.5 | Asn | 56 | 4.5 | 21 | 4 |
| 147.1 | 130 | 0.5 | Gln | 61 | 6 | 13 | 4 |
| 205.2 | 188.1 | 3.2 | Trp | 56 | 4.5 | 13 | 6 |
| 147.1 | 84.1 | 0.4 | Lys | 61 | 6 | 23 | 4 |
| 94.1 | 47.1 | 0.5 | U$^{13}$C,$^{15}$N-Ala | 51 | 5.5 | 17 | 4 |
| 110 | 63.1 | 0.5 | U$^{13}$C,$^{15}$N-Ser | 61 | 4.5 | 15 | 4 |
| 122.1 | 75 | 0.5 | U$^{13}$C,$^{15}$N-Pro | 61 | 7.5 | 19 | 4 |
| 124.1 | 77.2 | 0.5 | U$^{13}$C,$^{15}$N-Val | 56 | 4.5 | 15 | 4 |
| 125.1 | 78.2 | 0.5 | U$^{13}$C,$^{15}$N-Thr | 56 | 5 | 13 | 4 |
| 139.2 | 92.1 | 1.3 | U$^{13}$C,$^{15}$N-Ile + U$^{13}$C,$^{15}$N-Leu | 56 | 4.5 | 13 | 4 |
| 139.1 | 77.1 | 0.5 | U$^{13}$C,$^{15}$N-Asp | 56 | 10 | 19 | 4 |
| 154.1 | 107.1 | 0.5 | U$^{13}$C,$^{15}$N-Glu | 56 | 5.5 | 15 | 4 |
| 156.2 | 109.1 | 0.7 | U$^{13}$C,$^{15}$N-Met | 51 | 4 | 13 | 4 |
| 165.2 | 118.1 | 0.4 | U$^{13}$C,$^{15}$N-His | 61 | 5.5 | 17 | 4 |
| 176.2 | 129.2 | 2.6 | U$^{13}$C,$^{15}$N-Phe | 56 | 6 | 17 | 4 |
| 185.1 | 75.1 | 0.4 | U$^{13}$C,$^{15}$N-Arg | 66 | 6 | 31 | 4 |
| 192.1 | 145.2 | 1.4 | U$^{13}$C,$^{15}$N-Tyr | 56 | 7 | 17 | 4 |
| 154.1 | 136 | 0.5 | U$^{13}$C,$^{15}$N-Gln | 61 | 6 | 13 | 4 |
| 155.1 | 90.1 | 0.4 | U$^{13}$C,$^{15}$N-Lys | 61 | 6 | 23 | 4 |
| 210 | 193 | 2.8 | D$_5$-Trp | 20 | 4.5 | 13 | 6 |
| 210 | 192 | 2.8 | D$_5$-Trp-210-192 | 20 | 4.5 | 13 | 6 |
| 210 | 191 | 2.8 | D$_5$-Trp-210-191 | 20 | 4.5 | 13 | 6 |

**Supplemental Table S3.** Annotations of the MONA-Database - external provided in .xlsx-File

All processed datasets, including metabolite abundance matrices, and metadata required to reproduce the analysis, are archived on Zenodo https://doi.org/10.5281/zenodo.20049138.

### Supplemental Figures

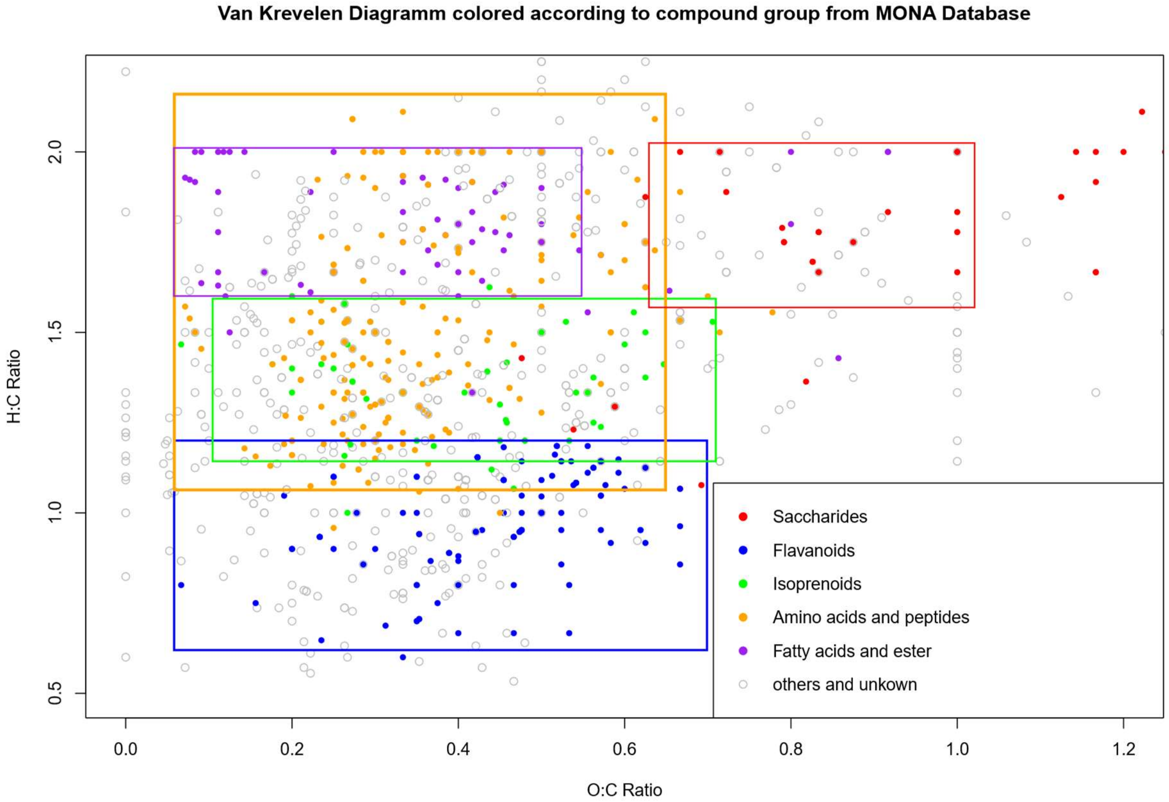


**Supplemental Figure S1. Van Krevelen-diagram coloured according to MONA-Annotation**

Van Krevelen-diagramm colored according to compound group from MassBank of North America MoNA Database (Company). The boxes include the theoretical H:C/O:C-Ratio for the named group, showing a good match between the theoretical sum formular and the annotated compounds.

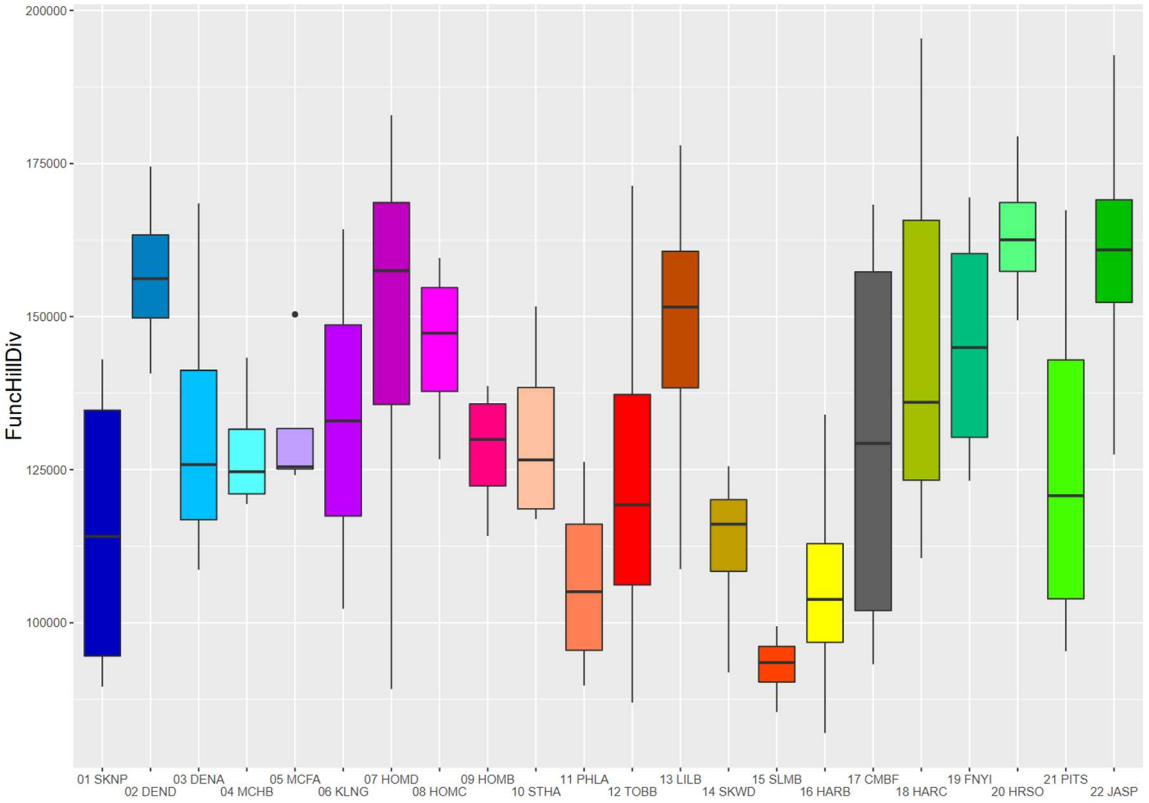


**Supplemental Figure S2. Functional-Hill Diversity between origins**

Functional Hill Diversity calculated for each individual tree and displayed as boxplots between different provenances from north (left) to south (right). The colour-coding matches the one in Figure 1. Calculations were done using the R-Package *chemodiv*.

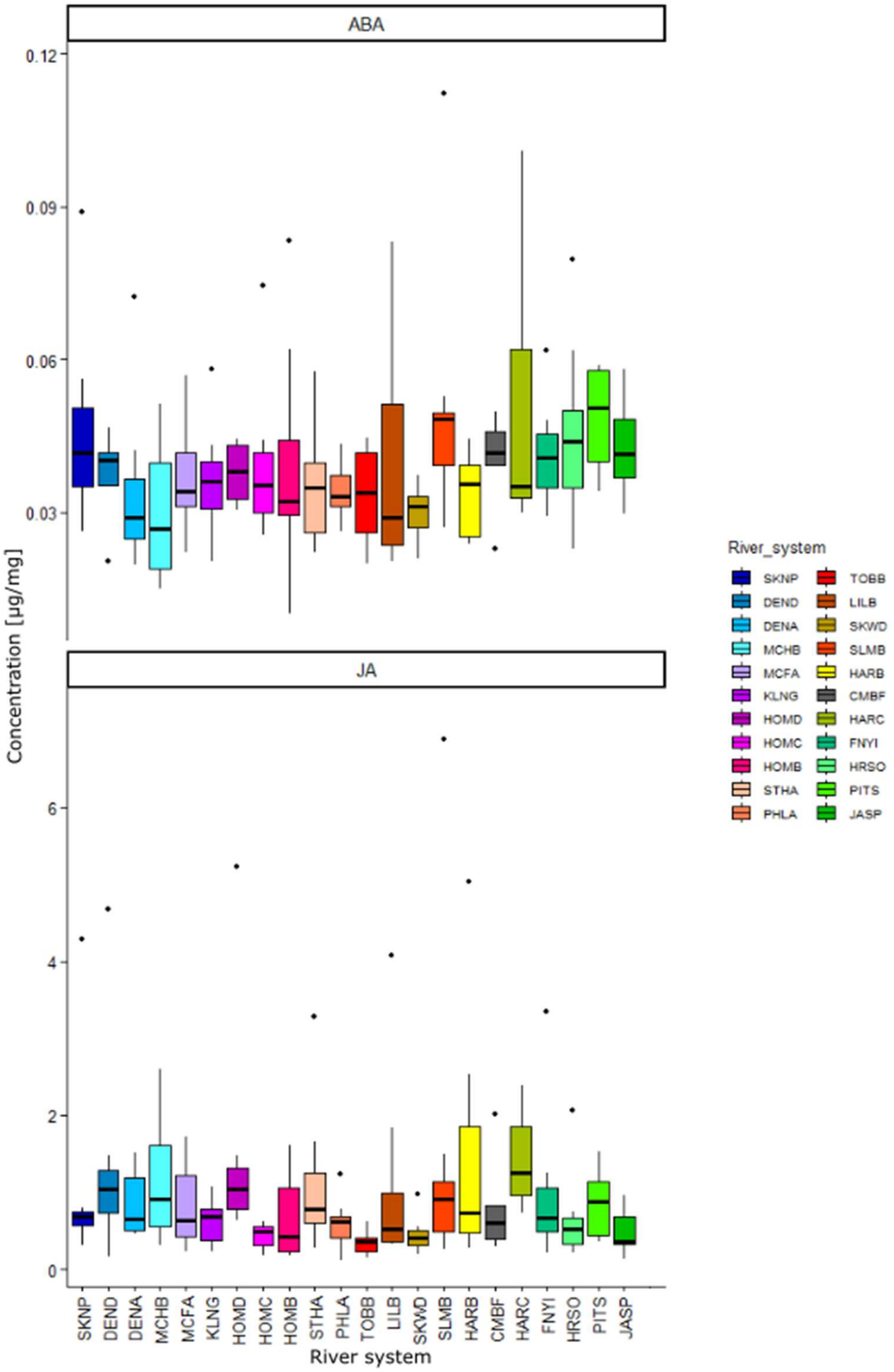


**Supplemental Figure S3 Additional phythormones between origins**

Boxplots for phytohormones over provenances: Jasmonic acid (JA) and abisic acid (ABA). All showing differences between provenances, but no trend can be observed. Provenances sorted from north (left) to south (right).

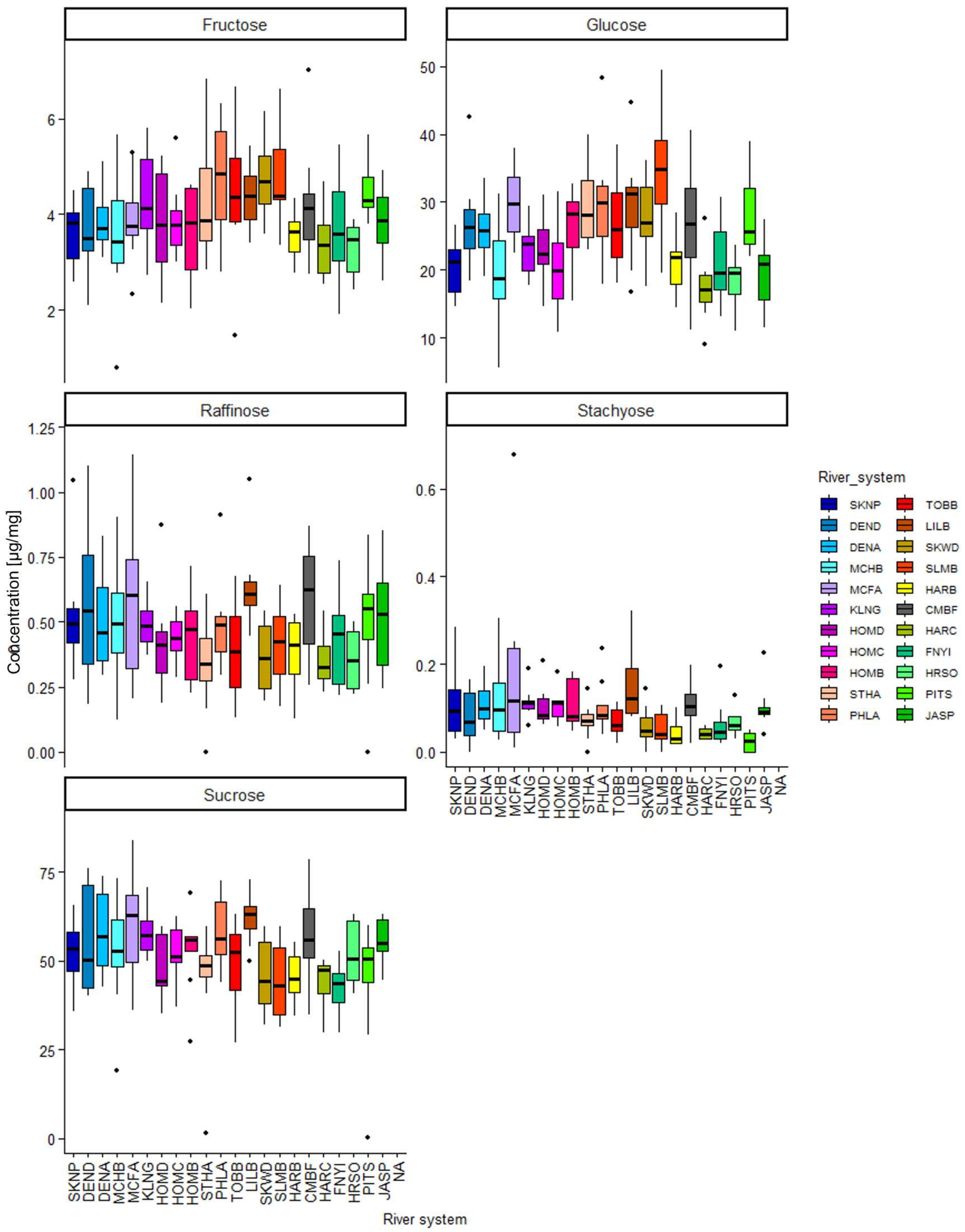


**Supplemental Figure S4. Sugars compounds between origins**

Boxplots for sugars over provenances: Fructose, glucose, raffinose, stachyose and sucrose All showing differences between provenances, but no trend can be observed. Provenances sorted from north (left) to south (right).

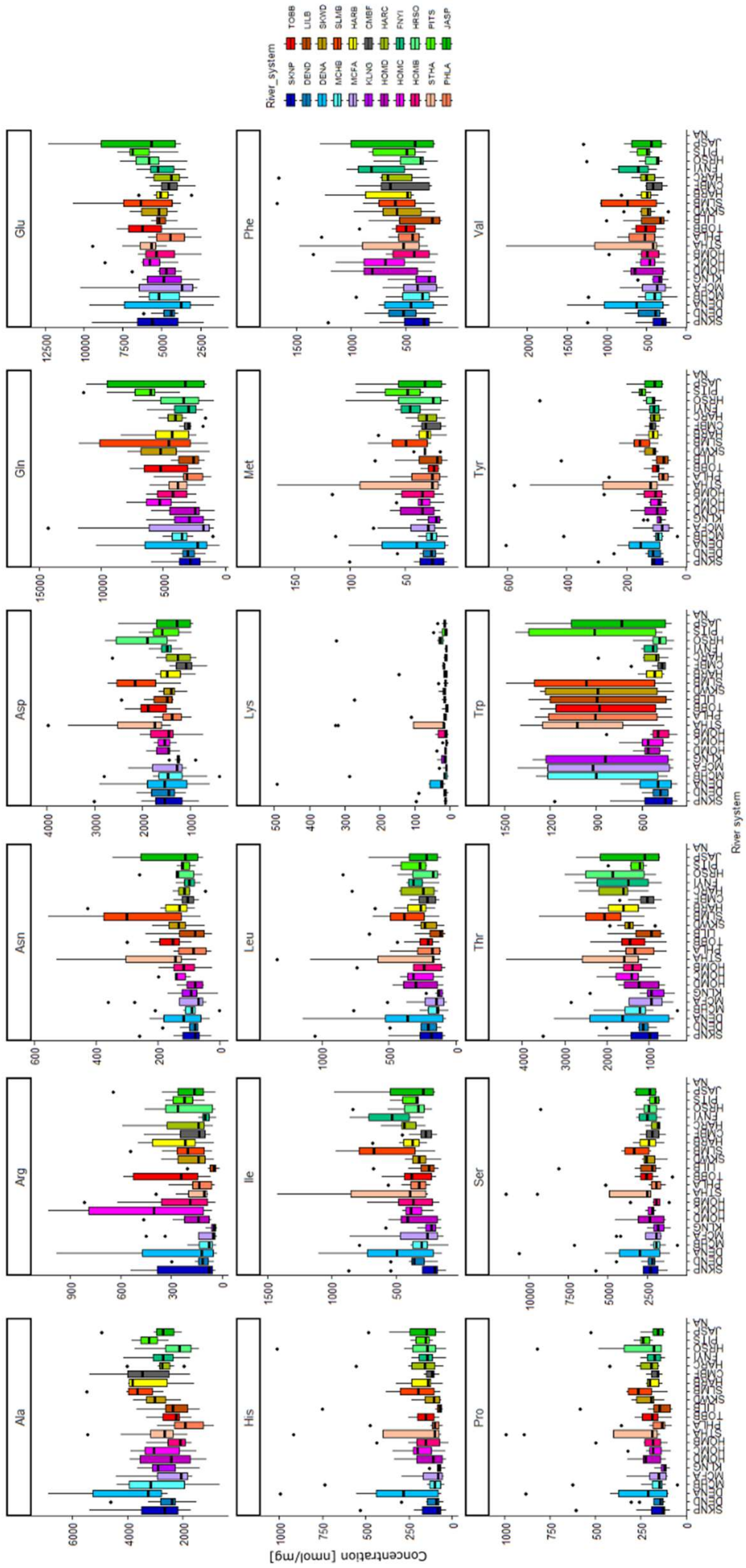


**Supplemental Figure S5. Amino acids between origins**

Boxplots for amino acids over provenances: Named with their three-letter abbreviation. All showing differences between provenances, but no trend can be observed. Provenances sorted from north (left) to south (right).

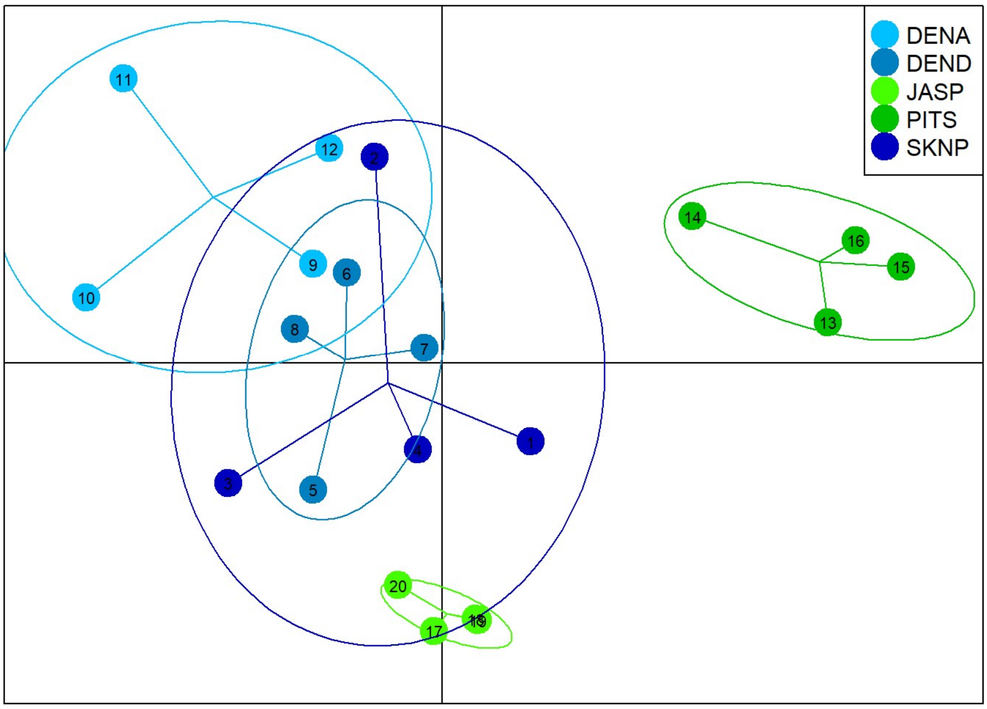


**Supplemental Figure S6. DAPC for targeted compounds for the North-South extremes according to Figure 5**

DAPC of the two most southern (JASP, PITS) and three most northern (SKNP, DENA, DEND) provenances using the targeted metabolomic data. Circles representing 90% confident intervals. Along the x-axis PITS and northern origins are well separated, while along the y-axis provenances PITS and northern origins are separated. Overall the separation is weaker than for the untargeted metabolomic data in Fig. 5.

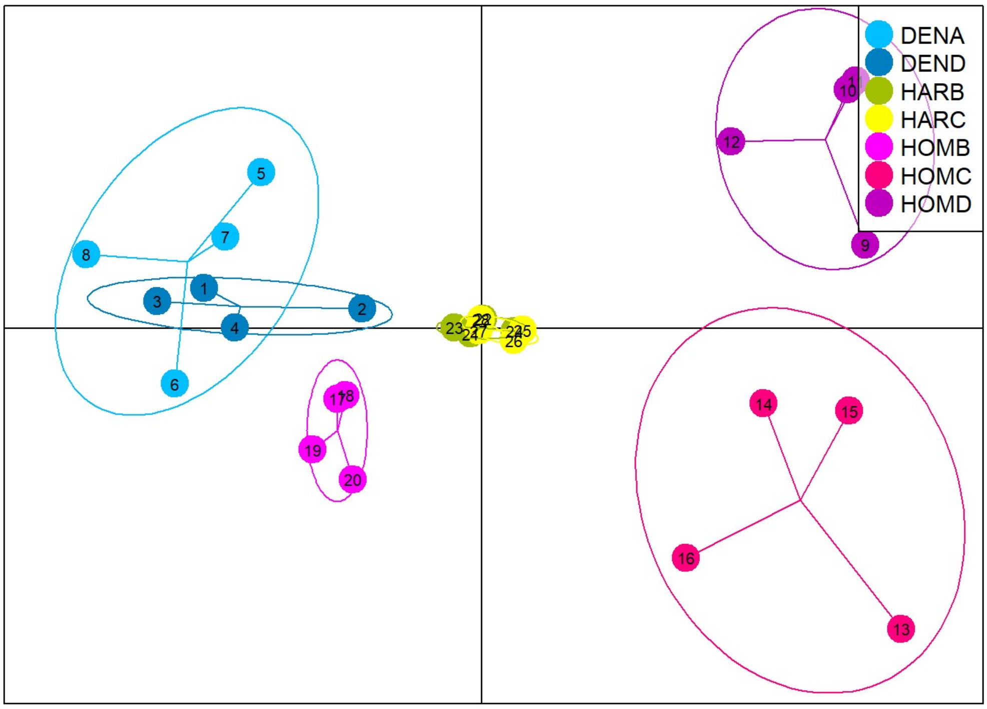


**Supplemental Figure S7. DAPC for targeted compounds for the North-South extremes according to Figure 6**

DAPC of of the three river systems (Homatoka, Dean, Harrison River)  using the targeted metabolomic data. Circles representing 90% confident intervals. There is a clear separation between the Harrison River (HAR) and the Dean River (DEN) but no clear separation between these two and the Homatoka River (HOM). Overall the separation is weaker than for the untargeted metabolomic data in Fig. 6.
